# Quantitative analysis of cis-regulatory elements in transcription with KAS-ATAC-seq

**DOI:** 10.1101/2024.02.29.582869

**Authors:** Ruitu Lyu, Yun Gao, Tong Wu, Chang Ye, Pingluan Wang, Chuan He

## Abstract

Cis-regulatory elements (CREs) are pivotal in orchestrating gene expression throughout diverse biological systems. Accurate identification and in-depth characterization of functional CREs are crucial for decoding gene regulation network and dynamics during cellular processes. In this study, we developed Kethoxal-Assisted Single-stranded DNA Assay for Transposase-Accessible Chromatin with Sequencing (KAS-ATAC-seq) to provide quantitative insights into transcriptional activity of CREs. A main advantage of KAS-ATAC-seq lies in its precise measurement of ssDNA levels within both proximal and distal ATAC-seq peaks, enabling the identification of transcriptional regulatory sequences in genomes. This feature is particularly adept at defining Single-Stranded Transcribing Enhancers (SSTEs). SSTEs are highly enriched with nascent RNA transcription and specific transcription factors (TFs) binding sites that determine cellular identity. Moreover, KAS-ATAC-seq provides a detailed characterization and functional implications of various SSTE subtypes; KAS-ATAC-seq signals on SSTEs exhibit more robust correlation with enhancer activities when compared with ATAC-seq data and active histone mark profiles. Our analysis of promoters and SSTEs during mouse neural differentiation demonstrates that KAS-ATAC-seq can effectively identify immediate-early activated CREs in response to retinoic acid (RA) treatment. We further discovered that ETS TFs and YY1 are critical in initiating early neural differentiation from mESCs to NPCs. Our findings indicate that KAS-ATAC-seq provides more precise annotation of functional CREs in transcription. Future applications of KAS-ATAC-seq would help elucidate the intricate dynamics of gene regulation in diverse biological processes and biomedical applications.

## Main

Gene expression regulation is largely mediated by cis-regulatory elements (CREs), which play a critical role in modulating gene functions across various biological processes^1–3^. CREs generally contain specific binding sites of transcription factors (TFs)^1^. DNA segments bound by TFs are often depleted of nucleosomes and are flanked by active histone marks^4^. Distal CREs, notably enhancers, engage in physical interactions with their target promoters, sometimes in a multilateral fashion^5–8^. Despite their crucial roles in creating cell type-specific transcriptomes, the precise mechanisms underlying the dynamic activation and precise looping of CREs have not been fully elucidated.

Distal CREs can be transcribed into both stable or unstable RNA transcripts, known as enhancer RNAs (eRNAs)^9,10^. Only a subset of distal CREs is capable of enhancing gene transcription^11^. Transcribed CREs typically demonstrate a stronger correlation with transcription activation and possess more functional relevance than those identified solely based on active histone marks, accessible chromatin regions, or DNase I hypersensitive sites (DHSs)^12–19^. The precise identification and in-depth characterization of these transcribed CREs are critical, especially in the context of cell differentiation, where subtle variations in gene regulation can result in significant phenotypic differences^20^. A variety of nascent RNA based approaches have been developed to study the dynamics of CREs transcription, including global run-on sequencing (GRO-seq)^21,22^, precision run-on sequencing (PRO-seq) and its variant PRO-CAP^23,24^, cap analysis of gene expression (CAGE) and NET-CAGE^25,26^, metabolic labeling with 4-thiouridine (4sU RNA)^27^, and mammalian native elongating transcript sequencing (mNET-seq)^28^. In addition, ATAC-seq has been developed for profiling chromatin accessibility^29^, and KAS-seq has been recognized for its rapid and sensitive detection of genome-wide single-stranded DNA (ssDNA) produced by transcriptionally active RNA polymerases in situ^30,31^, both serve as proxy of CRE activities.

These genomic methods all fill critical gaps, yet they also have limitations in terms of accuracy, sensitivity, or input material requirements. Nascent RNA-based methods are capable of directly detecting enhancer RNA (eRNA) but lack sufficient sensitivity due to the inherent instability of eRNA. Additionally, these methods are ineffective with limited starting materials. ATAC-seq and active histone marks are instrumental in defining enhancers, but they frequently can’t reflect the transcriptional activity, and a substantial number of distal CREs also function as poised enhancers and insulators. KAS-seq offers a promising alternative by efficiently detecting ssDNA on enhancers and gene transcription units, which is indicative of active transcription. However, it faces its own set of challenges, particularly in distinguishing enhancer-associated ssDNA from other ssDNA signals across the genome. These limitations highlight the need for a more refined approach that can overcome these shortcomings of existing methods, ensuring a more accurate and comprehensive understanding of enhancer dynamics and their impact on gene regulation.

In this study, we first developed an optimized KAS-seq (*Opti-*KAS-seq) protocol that significantly enhances the efficiency of capturing ssDNA. *Opti-*KAS-seq offers broader genomic coverage and higher signal-to-background ratio that works across a wide range of applications and sample types. By integrating the sensitive *Opti-*KAS-seq with ATAC-seq, we further introduce Kethoxal-Assisted Single-stranded DNA Assay for Transposase-Accessible Chromatin with Sequencing (KAS-ATAC-seq) with the dual capability to simultaneously uncover chromatin accessibility and transcriptional activity of CREs. A major advantage of KAS-ATAC-seq lies in its precise measurement of ssDNA levels within CREs, enabling the de novo identification of ssDNA promoter and Single-Stranded Transcribing Enhancers (SSTEs) as a subset of CREs without relying on eRNA or active histone marks ChIP-seq data. Additionally, we applied KAS-ATAC-seq to examine the transcriptional dynamics of CREs during the neural differentiation of mESCs into neural progenitor cells (NPCs). This analysis uncovered the involvement of specific transcription factors (TFs), including ETS and YY1, in the regulation of immediate-early activated promoters and SSTEs in response to RA treatment. These findings demonstrate the capability of KAS-ATAC-seq as a new powerful genomic method for precisely exploring and understanding the gene regulatory mechanisms by CREs.

## Results

### Enhancing ssDNA capture efficiency with optimized KAS-seq procedure

Transcription is a multifaceted and dynamic process that generates single-stranded DNA (ssDNA) regions in the genome, commonly referred as ’transcription bubbles’^32^. In our previous work, we developed KAS-seq to map transcriptional activities by sensitively capturing and sequencing genome-wide ssDNA through the N_3_-kethoxal–assisted labeling. Although current KAS-seq approach has proven to be effective in many contexts^33–38^, we and others have noticed compromised sensitivity of KAS-seq when using certain tissue samples and primary cells obtained using fluorescence-activated cell sorting (FACS). Our investigations suggested that this compromised efficiency in ssDNA capture might be due to the limited diffusion of N_3_-kethoxal through the cell membrane of these primary cells and tissues. We therefore modified the cell labeling procedure of KAS-seq by adding a cell permeabilization step, which allows N_3_-kethoxal to enter cells and label ssDNA more efficiently (Fig. 1a).

**Fig. 1.**
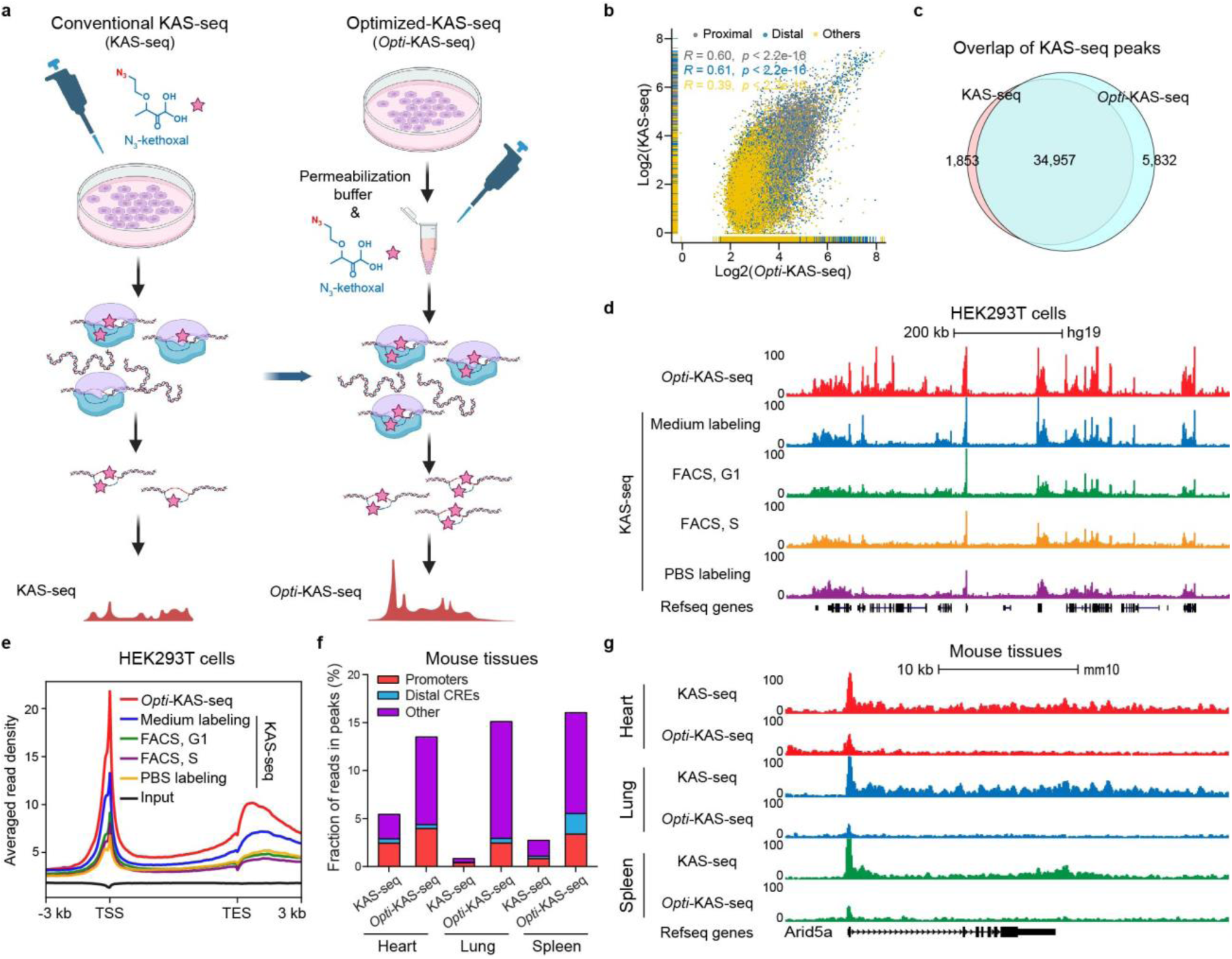
Optimization of the conventional KAS-seq protocol. **a**, Schematic comparing conventional KAS-seq (KAS-seq) and optimized KAS-seq (*Opti-*KAS-seq) protocols. KAS-seq involves N_3_-kethoxal labeling of cells directly in the culture dish medium, whereas Opti-KAS-seq first harvests and permeabilizes the cells, then performs N_3_-kethoxal labeling in a 1.5ml centrifuge tube. **b**, Scatterplot comparing KAS-seq and *Opti-*KAS-seq data in HEK293T cells across 1kb genomic bins on merged KAS-seq and *Opti-*KAS-seq peaks. Purple dots denote proximal genomic bins (n=13,308), blue dots denote distal genomic bins (n=6,609), and yellow dots denote other genomic bins (as labeled) (n=36467). **c**, Venn diagram illustrating the overlap of peaks identified using KAS-seq and *Opti-*KAS-seq data in HEK293T cells. **d**, Snapshot of UCSC genome browser tracks displaying the KAS-seq data generated using the *Opti-*KAS-seq and conventional KAS-seq methods for different types of applications across a representative region (chr1:23,572,909-24,358,036), including labeling of HEK293T cells directly in the culture dish medium (medium labeling), labeling of FACS-sorted cells at G1 (FACS, G1) and S (FACS, S) cell cycle in PBS, and labeling of harvested cells in PBS (PBS labeling). **e**, Metagene profile showing the distribution of KAS-seq signals from different protocols at gene-coding regions (n=36,231) in HEK293T cells, with 3 kb upstream of TSS and 3 kb downstream of TES shown. **f**, Stacked bar plot showing the proportion of reads in peaks that map to promoters (±500 bp from TSS), distal cis-regulatory elements (CREs) (>500 bp from TSS) and other regions from KAS-seq and *Opti-*KAS-seq datasets obtained from mouse heart, lung, and spleen tissues. These values were calculated based on 30 million randomly aligned de-duplicated reads. **g**. Snapshot of UCSC genome browser tracks displaying KAS-seq and *Opti-*KAS-seq datasets generated in mouse heart, lung, and spleen tissues across a representative region (chr1:36,306,107-36,331,699).

To confirm the effectiveness of the optimized KAS-seq (*Opti*-KAS-seq) protocol, we first tested it with HEK293T cells and conducted a thorough comparison between KAS-seq and *Opti-*KAS-seq under an equal number of uniquely mapped reads. Our quality control assessment revealed that the reproducibility, consistency, and robustness of *Opti-*KAS-seq match those of the conventional KAS-seq protocol (Extended Data Fig. 1a-f). A detailed exploration of KAS-seq peaks in HEK293T cells indicated that *Opti-*KAS-seq substantially elevates ssDNA detection sensitivity across promoters, distal enhancers, and other genomic regions (Fig. 1b-d). Comparative analysis between *Opti-*KAS-seq and KAS-seq, including peak overlaps (Fig. 1c), fingerprint plots (Extended Data Fig. 1g), and gene-coding enrichment (Fig. 1e), confirmed the expanded genomic coverage and elevated signal intensity achieved by *Opti-*KAS-seq (Fig 1d and Extended Data Fig. 1h-i). Moreover, *Opti-*KAS-seq mapped a larger fraction of sequencing reads to promoters, distal elements, and other genomic features than KAS-seq (Extended Data Fig. 1j). To validate our findings from HEK293T cells, we extended our analyses to E14-mESCs. The results consistently demonstrated the superior efficacy of *Opti-*KAS-seq in capturing ssDNA across the genome (Extended Data Fig. 2). We next applied *Opti-*KAS-seq to a variety of mouse tissues, including mouse heart, lung, and spleen, which were challenging for conventional KAS-seq. In these tissues, *Opti-*KAS-seq exhibited exceptional ssDNA capture efficiency (Fig. 1f-g and Extended Data Fig. 3). Taken together, these results highlight the advantages of *Opti-*KAS-seq over the conventional KAS-seq, particularly in improving ssDNA capture efficiency and expanding its applicability to previously challenging sample types.

### KAS-ATAC-seq simultaneously reveals chromatin accessibility and transcriptional activity of CREs

ATAC-seq detects accessible chromatin loci but it does not reveal transcription activity^39,40^. We envisioned that integrating ATAC-seq with KAS-seq would enable us to selectively capture ssDNA from CREs, thereby reflecting active transcription. This strategy aims to streamline the categorization of CREs and exclude ssDNA signals associated with non-regulatory regions commonly observed in KAS-seq data^30^. Our previous attempts have led to shallow peaks, likely due to reduced signal intensities^30^. By taking advantage of the enhanced ssDNA capture activity of *Opti*-KAS-seq, we further developed KAS-ATAC-seq that enables a comprehensive assessment of transcriptional activity of CREs in accessible chromatin regions. The integration of optimized N_3_-kethoxal-assisted ssDNA labeling with Tn5 transposase-mediated accessible chromatin detection enables intricate probing of transcriptional activities within CREs by capturing ssDNA at ATAC-seq peaks (Fig. 2a). The tagmentation step also simplifies library construction and allows application of KAS-ATAC-seq to limited input samples.

**Fig. 2.**
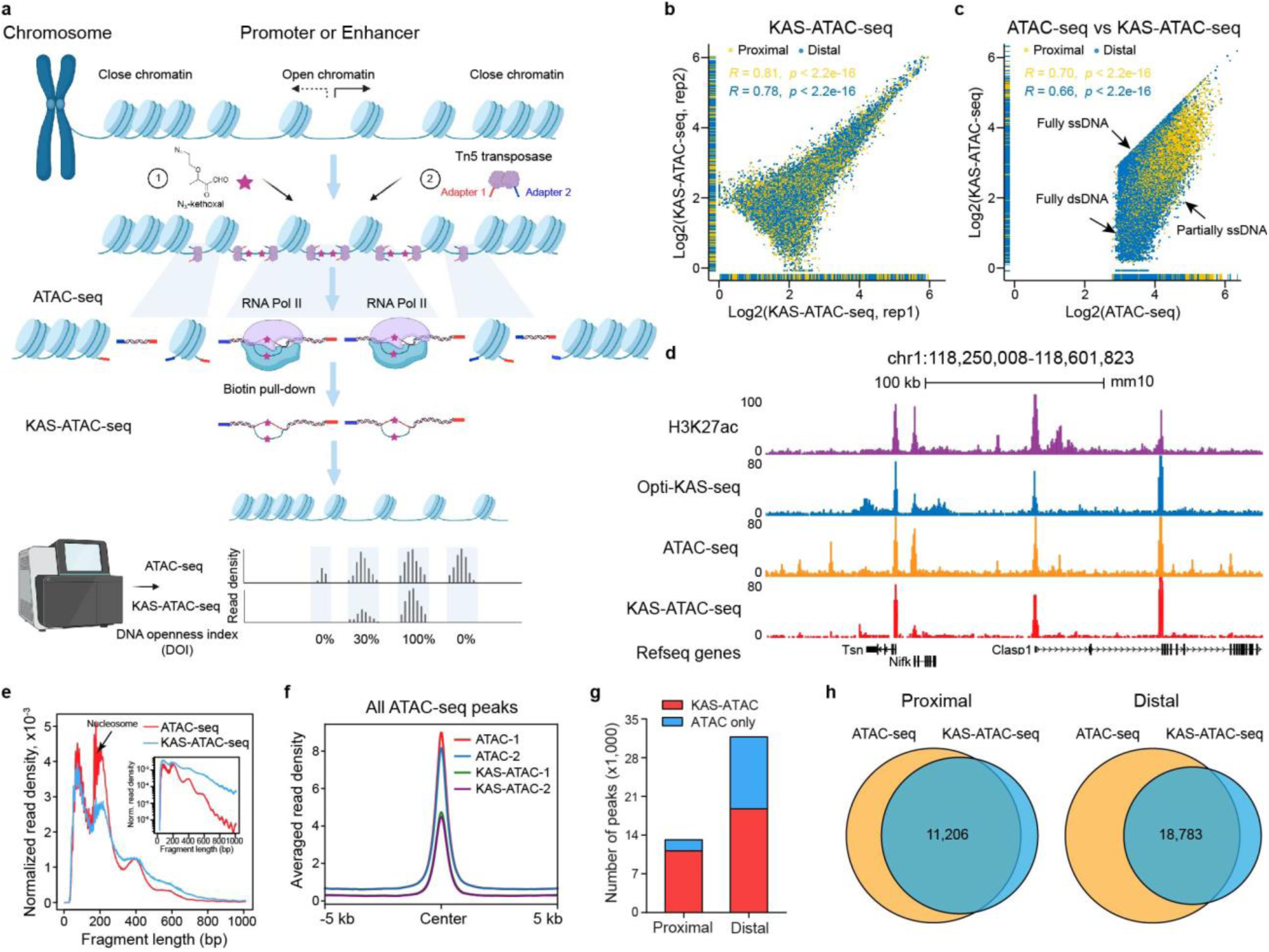
Development of KAS-ATAC-seq. **a**, Schematic of library preparation workflow in the KAS-ATAC-seq protocol. **b**, Scatterplot illustrating the Pearson correlation between two replicates of KAS-ATAC-seq data for proximal (n=11,522) and distal (n=25,561) peaks in fresh mESCs. Yellow dots represent proximal peaks, blue dots represent distal peaks. Both the Pearson correlation coefficients and significance (*p* values) for proximal and distal peaks are provided. **c**, Scatterplot illustrating the Pearson correlation between ATAC-seq and KAS-ATAC-seq data on proximal (n=13,158) and distal ATAC-seq peaks (n=31,768) in fresh mESCs. Yellow dots represent proximal ATAC-seq peaks, blue dots represent distal ATAC-seq peaks. Both the Pearson correlation coefficients and significance (*p* values) for proximal and distal peaks are provided. Peaks exhibiting higher signals in KAS-ATAC-seq relative to ATAC-seq are normalized to the ATAC-seq signals. **d**, Snapshot of UCSC genome browser tracks displaying H3K27ac ChIP-seq, *Opti-*KAS-seq, ATAC-seq, and KAS-ATAC-seq data on a representative region (chr1:118,250,008-118,601,823) featuring both proximal and distal KAS-ATAC-seq peaks. **e**, Density plot comparing the fragment sizes of ATAC-seq and KAS-ATAC-seq data generated in fresh mESCs. The inset, a log-transformed histogram, shows clear periodicity persists to nucleosomes. **f**, Metagene profile showing the distribution of averaged ATAC-seq and KAS-ATAC-seq read density across all ATAC-seq peaks identified in mESCs, with 5 kb upstream and 5 kb downstream from the center of ATAC-seq peaks shown. **g**, Stacked bar plot showing the numbers of ATAC-seq peaks, depicted by their overlap with KAS-ATAC-seq peaks in mESCs. Proximal and distal ATAC-seq peaks are displayed separately. **h**, Venn diagrams showing the overlap between ATAC-seq and KAS-ATAC-seq peaks identified in mESCs, with proximal and distal ATAC-seq and KAS-ATAC-seq peaks displaying separately.

KAS-ATAC-seq in mouse embryonic stem cells (mESCs) demonstrated high reproducibility between replicates, particularly in the characterization of proximal (n=11,522, R=0.81, *p*<2.2e-16) and distal (n=25,561, R=0.78, *p*<2.2e-16) KAS-ATAC-seq peaks (Fig. 2b). Through comparative analysis with ATAC-seq, our scatterplot investigations uncovered a detailed landscape of CREs. The majority of KAS-ATAC-seq peaks closely mirrored those detected in ATAC-seq. Furthermore, certain ATAC-seq peaks notably lacked enrichment of ssDNA signals (Fig. 2c), emphasizing the refined specificity of KAS-ATAC-seq in identifying CREs in transcription. We observed three distinct patterns: (1) CREs showing consistent signal intensities across both methods (fully ssDNA); (2) CREs with reduced KAS-ATAC-seq read densities compared to those of ATAC-seq (partially ssDNA at accessible CREs); and (3) CREs with complete absence of KAS-ATAC-seq peaks but clear ATAC-seq peaks (fully dsDNA at accessible CREs). These observations underscore the capability of KAS-ATAC-seq to offer a more nuanced perspective on transcription across CREs that are accessible based on ATAC-seq alone (Fig. 2d).

In our investigation of Tn5 transposase-accessible chromatin, we observed a marked difference in fragment size distribution between ATAC-seq and KAS-ATAC-seq libraries. Specifically, ATAC-seq captures more DNA fragments that contain mono-nucleosomes (∼200 bp), whereas KAS-ATAC-seq captures a greater proportion of ssDNA fragments in nucleosome-free regions (<100 bp), offering potential unique insights into the dynamics between transcription initiation and nucleosome positioning (Fig. 2e). KAS-ATAC-seq signals align mostly within the ATAC-seq spectrum, but with lower average read density across all ATAC-seq peaks in E14-mESCs (Fig. 2f). Additionally, a substantial portion (78.5%, 11,206/14,277) of proximal ATAC-seq peaks intersect with KAS-ATAC-seq peaks, in contrast to only 45.8% (18,783/41,007) for distal ATAC-seq peaks. This indicates that KAS-ATAC-seq captures a more pronounced ssDNA presence in proximal ATAC-seq peaks compared to distal ATAC-seq peaks (Fig. 2g-h). We further extended our KAS-ATAC-seq protocol to HEK293T cells0. The KAS-ATAC-seq read intensities across proximal and distal regions are consistent with the results in E14-mESCs, supporting the robustness of KAS-ATAC-seq across different cell types (Extended Data Fig. 4a-c). Similar patterns in mESCs were also observed in HEK293T cells when inspecting the genomic footprints and loci-specific interactions between KAS-ATAC-seq and ATAC-seq (Extended Data Fig. 4d-e). Therefore, we establish KAS-ATAC-seq as a tool ideally suited to elucidate transcriptional activities through ssDNA capture within CREs delineated by ATAC-seq peaks.

### Quantitative analysis of CRE activity using the DNA Openness Index (DOI)

We devised the DNA Openness Index (DOI), a novel metric specifically designed to evaluate the openness of double-stranded DNA (dsDNA). This is achieved by calculating the ratio of KAS-ATAC-seq to ATAC-seq signals across both proximal and distal CREs. The DOI thus offers insights into the transcriptional activity within these regulatory sequences, serving as a quantitative indicator of DNA transcriptional engagement (Fig. 3a). Interestingly, a higher proportion of distal CREs (19.3%, 7,897/41,007) were observed as fully ssDNA (DOI:100%) in comparison to proximal CREs (12.2%,1,736/14,277) (Fig. 3a). In the meantime, proximal CREs typically display elevated DOI values relative to distal CREs in E14-mESCs, with this difference being particularly notable in partially ssDNA CREs (Fig. 3b). This implies a more active role for proximal CREs in regulating gene expression compared to distal CREs in E14-mESCs, possibly due to their closer proximity to transcription start site (TSS). DOI values for both proximal and distal CREs remained consistent across different chromosomes (Extended Data Fig. 5a). Categorizing CREs in E14-mESCs based on their DOI values (high, medium, low, and zero) revealed that the variation of DOI is primarily dependent on KAS-ATAC-seq signals rather than ATAC-seq signals (Fig. 3c). This observation suggests that CREs with comparable levels of chromatin accessibility can exhibit markedly different transcriptional activities. Additionally, we noticed an abundance of CpG-rich sequence motifs specifically enriched on CREs with high-DOI values (Fig. 3d). This pattern is further supported by the finding that CREs with high-DOI values generally exhibit a higher density of CpG sites (Fig. 3e).

**Fig. 3.**
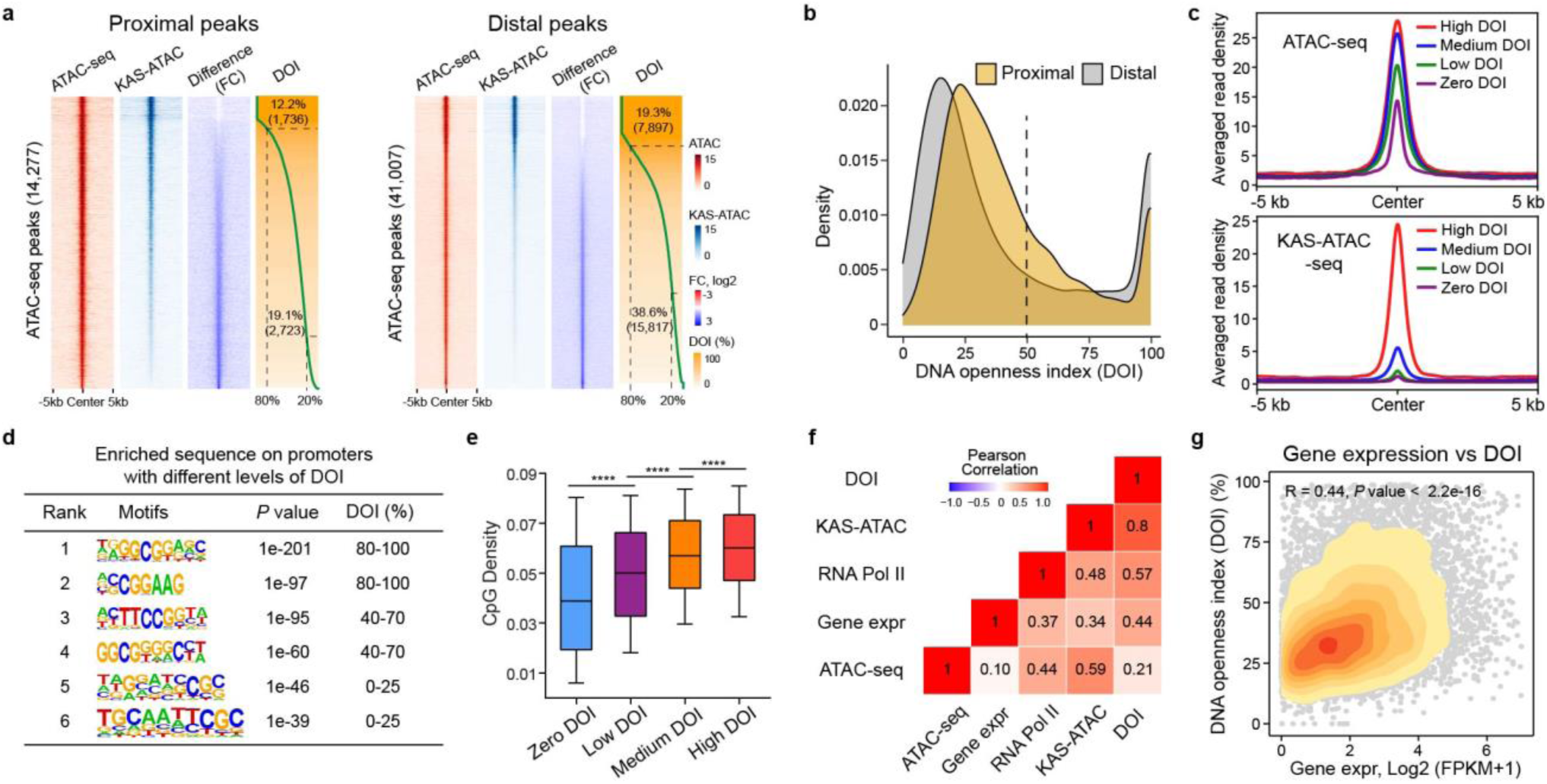
KAS-ATAC-seq quantitatively estimates the dsDNA openness in ATAC-seq peaks. **a**, Heatmap showing the ATAC-seq, KAS-ATAC-seq, log2 fold changes between KAS-ATAC-seq and ATAC-seq, and the DNA openness index (DOI) for proximal (left panel, n=14,277) and distal (right panel, n=41,007) ATAC-seq peaks. The DOI is determined as the proportion of KAS-ATAC-seq signal relative to the ATAC-seq signal at a specific CRE. Regions spanning 5 kb upstream and 5 kb downstream from the center of proximal and distal ATAC-seq peak were shown. Cis-regulatory elements (CREs) with DOI values exceeding 80% or below 20% are demarcated with dashed lines in the DOI heatmap. **b**, Density plot comparing the DNA openness index (DOI) of proximal (n=14,277) and distal (n=41,007) ATAC-seq peaks. **c**, Metagene profiles showing the distribution of averaged ATAC-seq (upper panel) and KAS-ATAC-seq (lower panel) read density across ATAC-seq peaks categorized by high, medium, low, zero DNA openness index (DOI) in mESCs. Regions spanning 5 kb upstream and 5 kb downstream from the center of ATAC-seq peaks are displayed. **d**, Table presenting the enriched sequence features on cis-regulatory elements (CREs) associated with varying levels of the DNA openness index (DOI). **e**, Boxplot illustrating the CpG density within proximal ATAC-seq peaks, categorized by high (n=4,386), medium (n=4,386), low (n=4,386), and zero (n=2,001) DNA openness index (DOI) in mESCs. The box shows 1st quartile, median and 3rd quartile, respectively. **f**, Correlation heatmap showing Pearson correlation coefficients between the DNA openness index (DOI), ATAC-seq, KAS-ATAC-seq, RNA Pol II binding, and gene expression levels of genes with ATAC-seq peaks (n=10,185). The gene expression levels of proximal ATAC-seq peaks target genes are determined using bulk cell RNA-seq data. The Pearson correlation coefficients were labeled on the correlation heatmap. **g**, Scatterplot showing the Pearson correlation between DNA openness index (DOI) and gene expression (Gene expr, bulk cell RNA-seq) on genes with ATAC-seq peaks (n=10,185). The Pearson correlation coefficient (R) and its associated *p* value are displayed at the top of the plot. Points are color-coded in orange to indicate the gene density.

Our analysis also identified proteins exhibiting strong correlations with DOI, particularly in high-DOI CREs of E14-mESCs. These include the zinc finger protein (E2f1), components of the mediator complex (Med1), cyclin-dependent kinases (Cdk7), and other specific TFs such as TBP, Brd4, Chd2, and Taf3 (Extended Data Fig. 5b-c). This indicates that these TFs play a more important role in regulating CRE transcription compared to other TFs. In addition, we generated a correlation heatmap to depict the relationships among DOI, ATAC-seq, KAS-ATAC-seq, RNA Pol II binding, and gene transcription as determined by RNA sequencing. The heatmap clearly reveals a stronger correlation between DOI metrics or KAS-ATAC-seq with gene transcription, in contrast to the relatively weak correlation observed between ATAC-seq data or active histone marks (H3K27ac and H3K4me3) and gene transcription (Fig. 3f-g and Extended Data Fig. 5d-h). This highlights the enhanced predictive power of DOI and KAS-ATAC-seq for capturing gene expression dynamics compared to ATAC-seq. In summary, our study introduces the DOI as a new metric for evaluating dsDNA openness across CREs in transcription. Moreover, both KAS-ATAC-seq and DOI emerge as more accurate indicators of gene expression.

### De novo mapping of Single-Stranded Transcribing Enhancers (SSTEs) using KAS-ATAC-seq

Numerous studies have established that only a subset of active enhancers is capable of enhancing gene transcription, and this capability is closely associated with the level of eRNA present on transcribed enhancers^11,19^. Building upon our prior work that highlighted the effectiveness of KAS-seq in detecting transcriptionally active enhancers through profiling ssDNA produced by RNA polymerases^31^, we have advanced our methodology. Specifically, we have annotated a specific group of distal CREs using KAS-ATAC-seq without relying on nascent RNA-seq or active histone marks. These elements, termed as Single-Stranded Transcribing Enhancers (SSTEs) (Fig. 4a), are characterized as KAS-ATAC-seq peaks that are frequently associated with RNA Pol II binding (57.4%, 10,774/18,783) and exhibit RNA transcription (Fig. 4b). Our cumulative frequency analysis revealed that approximately 60% of SSTEs exhibit detectable eRNA transcripts in at least one of six different nascent RNA-seq assays from mESCs, including GRO-seq, PRO-seq, PRO-cap, NET-CAGE, mNET-seq, and 4sU RNA-seq, all targeting newly synthesized nascent RNA transcripts (Fig. 4c). In contrast, Double-Stranded Elements (DSEs), defined as ATAC-seq peaks not overlapping with KAS-ATAC-seq peaks, predominantly lack detectable ssDNA and eRNA transcripts (Fig. 4d-e and Extended Data Fig. 6a). Notably, KAS-ATAC-seq signals exhibit a stronger correlation with nascent RNA transcription than ATAC-seq signals on SSTEs (Extended Data Fig. 6b). These SSTEs revealed by KAS-ATAC-seq is therefore a more sensitive and straightforward way to define transcriptionally active enhancers compared to traditional nascent RNA-seq assays and ATAC-seq (Fig. 4g).

**Fig. 4.**
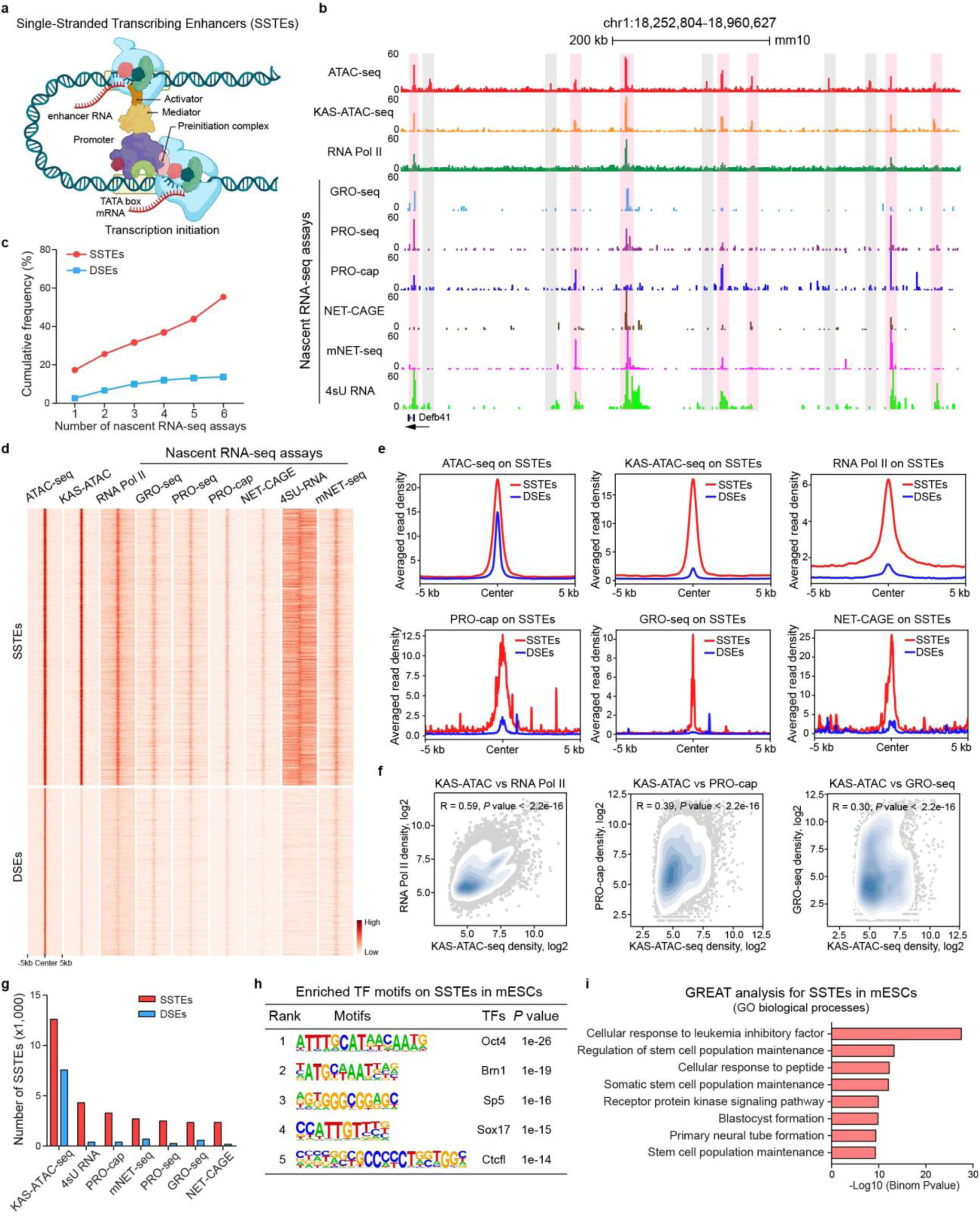
Prominent nascent RNA transcription is evident on Single-Stranded Transcribing Enhancers (SSTEs) identified by KAS-ATAC-seq. **a**, Schematic illustrating transcription initiation at promoters and enhancers, as well as the long-range interactions between them. **b**, Snapshot of UCSC genome browser tracks displaying ATAC-seq, KAS-ATAC-seq, RNA Pol II, and various nascent RNA-seq datasets (GRO-seq, PRO-seq, PRO-cap, NET-CAGE, mNET-seq, and 4sU RNA-seq) on a representative region (chr1:18,252,804-18,960,627) featuring cis-regulatory elements (CREs) defined by ATAC-seq and KAS-ATAC-seq peaks. SSTEs are defined as ATAC-seq peaks that overlap with KAS-ATAC-seq peaks and are highlighted in pink. DSEs are defined as ATAC-seq peaks without KAS-ATAC-seq overlap and are highlighted in grey. **c**, Line graph depicting the cumulative frequency of SSTEs (n=12,601) and DSEs (n= 7,574) with significant nascent RNA transcription detected by various nascent RNA-seq datasets (GRO-seq, PRO-seq, PRO-cap, NET-CAGE, mNET-seq, and 4sU RNA-seq). The red dotted line represents SSTEs, and the green dotted line represents DSEs. **d**, Heatmap showing the enrichment of ATAC-seq, KAS-ATAC-seq, RNA Pol II, and various nascent RNA transcription signals (GRO-seq, PRO-seq, PRO-cap, NET-CAGE, mNET-seq, and 4sU RNA-seq) on intergenic SSTEs (n=12,601, top panel) and DSEs (n= 7,574, bottom panel). Regions spanning 5 kb upstream and 5 kb downstream from the center of CREs are shown. **e**, Metagene profiles showing the averaged read density of various nascent RNA transcription datasets (GRO-seq, PRO-seq, PRO-cap, NET-CAGE, mNET-seq, and 4sU RNA-seq) across SSTEs (red) and DSEs (blue). **f**, Scatterplot showing the Pearson correlation between KAS-ATAC-seq data and RNA Pol II (left), PRO-cap (middle), and GRO-seq (right) data on intergenic SSTEs (n=12,601). The Pearson correlation coefficient (R) and its associated p value are displayed at the top of the plot. Points are color-coded in light blue to indicate SSTE density. **g**, Grouped bar plot displaying the number of SSTEs identified by KAS-ATAC-seq data alongside various nascent RNA transcription datasets, including GRO-seq, PRO-seq, PRO-cap, NET-CAGE, mNET-seq, and 4sU RNA-seq. **h**, Table presenting the enriched motifs of transcription factors (TFs) identified on SSTEs in mESCs. **i**, Horizontal bar plot illustrating the Gene Ontology (GO) biological processes derived from the GREAT analysis of SSTEs in mESCs.

Active histone marks are significantly abundant on SSTEs. Specifically, H3K4me3 and H3K36me3 are exclusively enriched on SSTEs. However, both SSTEs and DSEs exhibit substantial enrichment of H3K27ac and H3K4me1 (Extended Data Fig. 6c-e). STARR-seq (Self-Transcribing Active Regulatory Region sequencing) is a powerful technique developed to quantify enhancer activity across the genome. Interestingly, we found that KAS-ATAC-seq signals on SSTEs align more closely with STARR-seq data than ATAC-seq profiles (Extended Data Fig. 6f-g), indicating that KAS-ATAC-seq effectively identifies functional CREs and reflects their activities. Consensus sequence motif analysis revealed enrichment of specific transcription factors such as Oct4, Brn1, Sp5, and Sox17 on SSTEs (Fig. 4h). The GREAT analysis further revealed that SSTEs are closely associated with biological processes involved in stem cell maintenance in mESCs (Fig. 4i). Conversely, DSEs, despite being enriched with ATAC-seq peaks and active histone marks, are primarily linked to signal transduction and cell differentiation (Extended Data Fig. 6h). This distinction highlights the functional importance of SSTEs in determining stem cell identity of mESCs (Fig. 4i). Additionally, SSTEs exhibit lower evolutionary conservation scores compared to DSEs and promoters, suggesting a higher level of cell-type specificity and potentially greater sequence variability (Extended Data Fig. 6i-l). Collectively, KAS-ATAC-seq enables de novo annotation of SSTEs in transcription as a subset of distal CREs. These SSTEs are distinguished by active RNA transcription, unique chromatin features, and specific TFs binding with functional importance.

### Transcriptional dynamics of promoters and SSTEs during neural differentiation from mouse embryonic stem cells

To assess the effectiveness of KAS-ATAC-seq in characterizing CREs during a continuous differentiation process, we examined the transcriptional dynamics of CREs throughout the neural differentiation process from mESCs to embryoid bodies (EBs) and neural progenitor cells (NPCs). Following the established protocol by Xiang et al.^41^, we conducted an time-course analysis employing both ATAC-seq and KAS-ATAC-seq. The neural differentiation was initiated by the removal of leukemia inhibitory factor (LIF) at Day 0, followed by treatment with retinoic acid (RA) at Day 2, resulting in the formation of NPCs by Day 8 (Fig. 5a-b). Our analysis revealed a gradual deactivation of SSTEs associated with pluripotency genes (Oct4 and Nanog) and an activation of SSTEs near early neural marker genes (*Pax6* and *Neurog2*), confirming the reliability of our time-course ATAC-seq and KAS-ATAC-seq data (Extended Data Fig. 7a-b). Furthermore, we found that KAS-ATAC-seq profiles offer a more distinct separation than ATAC-seq profiles of the regulome across various stages of neural differentiation (Fig. 5c). This highlights notable changes in the transcriptional dynamics of promoters and SSTEs during the neural differentiation of mESCs into NPCs.

**Fig. 5.**
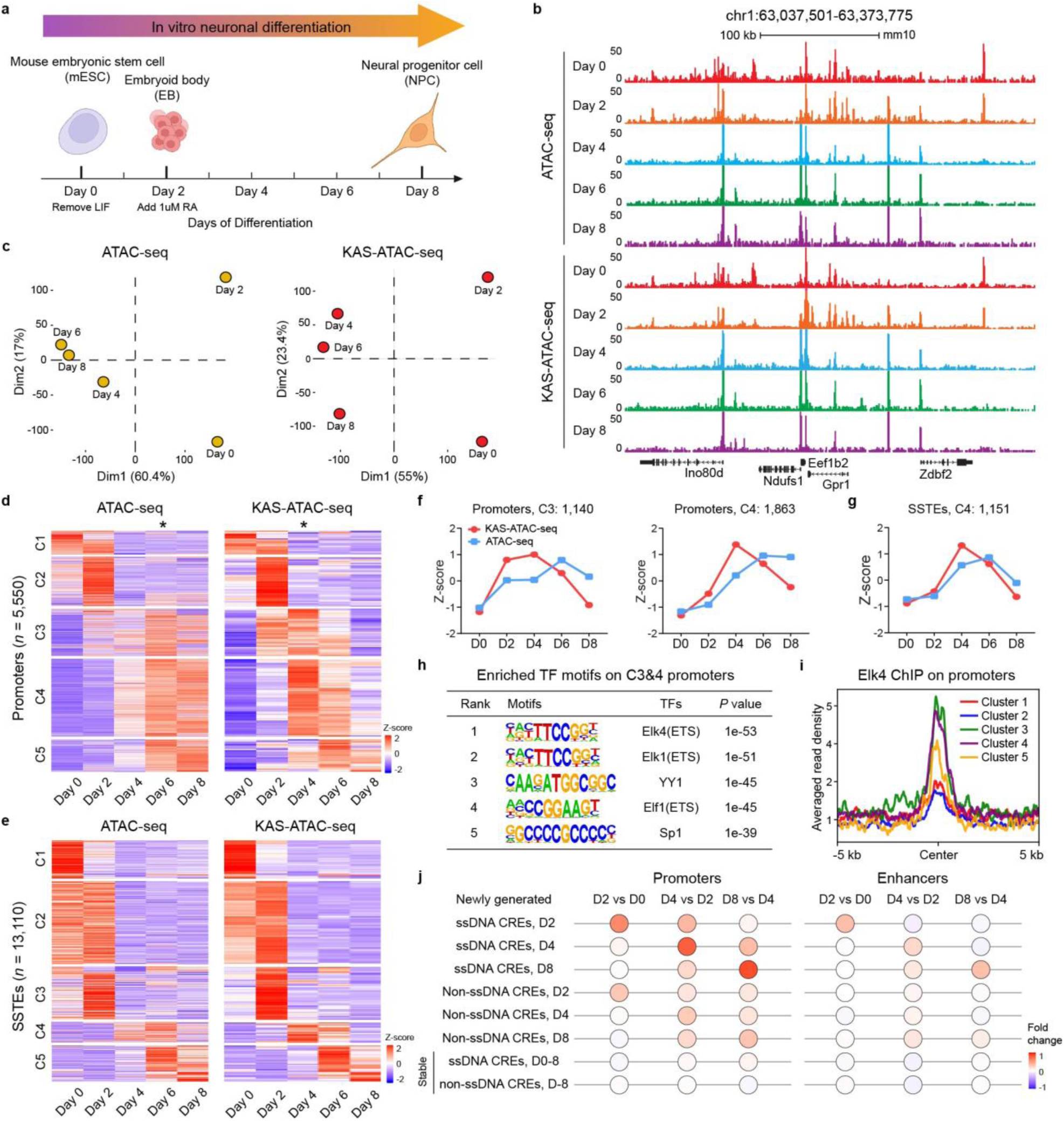
KAS-ATAC-seq accurately captures the transcriptional dynamics of ssDNA promoters and SSTEs during neural differentiation. **a**, Schematic of the in vitro neural differentiation procedure from mouse embryonic stem cells (mESCs) to neural progenitor cells (NPCs). mESCs were initially used to form embryoid bodies (EBs) by withdrawing leukemia inhibitory factor (LIF), followed by induction of the neural differentiation of EBs into NPCs using retinoic acid (RA) treatment. Two biologically independent samples were harvested at the indicated time points for ATAC-seq and KAS-ATAC-seq. **b**, Snapshot of UCSC genome browser tracks displaying the ATAC-seq and KAS-ATAC-seq data for a representative region (chr1:63,037,501-63,373,775). ATAC-seq and KAS-ATAC-seq data were generated using cells harvested at different time points indicated during the neural differentiation process. **c**, Principal component analysis (PCA) plot of ATAC-seq (left panel) and KAS-ATAC-seq (right panel) data generated using cells harvested at different time points indicated during the in vitro neural differentiation process. **d**,**e**, Heatmaps of time-course ATAC-seq and KAS-ATAC-seq profiles during the neural differentiation of mESCs into NPCs, using significantly dynamic CREs (up- and down-regulated), clustered on the basis of the KAS-ATAC-seq data. Each row represents a promoter (**d**) and SSTE (**e**) and each column represents a time point. The row order is the same between ATAC-seq (left panel) and KAS-ATAC-seq (right panel). The averaged values of two biologically independent samples at each time point are shown. Asterisks represent peak time points of promoters in the heatmaps of ATAC-seq and KAS-ATAC-seq data. **f**, Line graph depicting the z-scores of C3 (left panel, n=1,140) and C4 (right panel, n=1,863) promoters in the clustered heatmap of dynamic promoters (**d**). The z-scores were calculated using ATAC-seq data (blue) and KAS-ATAC-seq data (red) at various stages of the mouse neural differentiation. **g**, Line graph depicting the z-scores of C4 SSTEs (n=1,151) in the clustered heatmap of dynamic SSTEs (**e**). The z-scores were calculated using ATAC-seq data (blue) and KAS-ATAC-seq data (red) at various stages of the mouse neural differentiation. **h**, Table presenting the enriched motifs of transcription factors (TFs) identified on cluster 3&4 (C3&4) immediate-early activated promoters in the clustered heatmap (**d**) during the mouse neural differentiation. **i**, Metagene profile showing the averaged read density of Elk4 ChIP-seq data over different promoter clusters in the clustered heatmap (**d**) during the mouse neural differentiation. **j**, Bubble plots showing the fold changes of gene expression levels between consecutive stages for genes with newly generated and stable ssDNA and non-ssDNA CREs defined by KAS-ATAC-seq and ATAC-seq data during the mouse neural differentiation from mESCs to EBs and NPCs (D0 to D8). Newly generated CREs are shown separately as promoters and enhancers. The color key, ranging from blue to red, indicates the median of fold changes of gene expression levels from low to high, respectively. The bulk RNA-seq data are referred from GSE151900.

Utilizing KAS-ATAC-seq, we identified 5,550 ssDNA promoters and 13,110 SSTEs that undergo dynamic changes at least once during the process of neural differentiation (Extended Data Fig. 7c-d). GREAT analysis of these dynamic CREs revealed that up-regulated ssDNA promoters and SSTEs are predominantly associated with neurogenesis and nervous system development (Extended Data Fig. 7e), whereas down-regulated ssDNA promoters and SSTEs are linked to stem cell maintenance and embryonic pattern specification (Extended Data Fig. 7f). This distribution reflects the cell identity transition from mESCs to NPCs. In contrast, dynamic DSEs and non-ssDNA promoters, which showed significant changes in ATAC-seq signals but not in KAS-ATAC-seq signals (Extended Data Fig. 7g-h), did not exhibit these specific associations, underscoring the unique regulatory roles of dynamic SSTEs in guiding the neural differentiation pathway.

Intriguingly, we found that a subset of promoters (C3:1,140; C4:1,863) and SSTEs (C4: 1,151), up-regulated by retinoic acid (RA) treatment (Supplementary Table 1), exhibited an earlier peak activation in KAS-ATAC-seq at Day 4, compared to ATAC-seq at Day 6 (Fig. 5d-g and Extended Data Fig. 7i-j). This is likely due to the fact that the KAS-ATAC-seq signal reflects real-time transcription levels by detecting ssDNA, whereas the ATAC-seq signal indicates chromatin accessibility on CREs. Additionally, we found that the binding motifs of ETS subfamily TFs and YY1 are particularly abundant in these early activated CREs (Fig. 5h). The ChIP-seq analysis of Elk4 and YY1 confirmed their significant binding enrichment on both early activated ssDNA promoters and SSTEs compared to other CREs (Fig. 5i and Extended Data Fig. 7k-m), suggesting that ETS TFs and YY1 act as key drivers in initiating neural differentiation, especially during the critical period induced by RA treatment from Day 2 to Day 4. In summary, our findings underscore the capability of KAS-ATAC-seq to profile the transcriptional dynamics of promoters and SSTEs with higher temporal resolution than ATAC-seq during neural differentiation.

To investigate the temporal relationship between dynamic CREs and gene expression during mouse neural differentiation, we integrated our KAS-ATAC-seq and ATAC-seq profiles with existing RNA sequencing (RNA-seq) data from mESCs (D0) to EBs (D2) and NPCs (D8). We focused on genes with dynamic promoters and enhancers that show different ATAC-seq or KAS-ATAC-seq signals at early and late time points of neural differentiation. The normalized fold changes in expression levels of these genes were calculated across consecutive developmental stages (Fig. 5j and Extended Data Fig. 8a). We found that the expression level of genes with newly generated ssDNA promoters and SSTEs in EBs and NPCs were up-regulated in these stages (Fig. 5j). Conversely, genes losing ssDNA promoters and SSTEs in EBs and NPCs demonstrated down-regulation in gene expression within these stages (Extended Data Fig. 8a). Notably, ssDNA promoters displayed a more significant temporal relationship with gene expression compared to SSTEs. Genes associated with dynamic promoters and enhancers lacking ssDNA showed a weak temporal regulation pattern, while genes with stable promoters and enhancers without ssDNA changes exhibited no discernible temporal regulation pattern (Fig. 5j and Extended Data Fig. 8a). Furthermore, we observed that ssDNA promoters and SSTEs undergo distinct transitions during the later stages of neural differentiation, potentially transitioning into other CRE subtypes, including non-ssDNA promoters, DSEs, and beyond CRE classification (no CREs) (Extended Data Fig. 8b). By categorizing ssDNA promoters and SSTEs based on their transition into other CRE subtypes, we discovered that those which did not transition into other CRE types displayed significantly higher KAS-ATAC-seq signal intensities (Extended Data Fig. 8c). However, this significance is less pronounced in ATAC-seq and H3K27ac profiles (Extended Data Fig. 8d-e). This suggests that the presence of ssDNA could serve as a predictive marker for the stability of CREs during neural differentiation, highlighting the potential of KAS-ATAC-seq in providing insights into the dynamic regulatory landscape of cell differentiation.

### Characterization and functional implications of various SSTE subtypes

In our analysis of SSTEs, we observed that H3K27ac peaks in a subset of SSTEs notably extends beyond the boundaries of KAS-ATAC-seq and ATAC-seq peaks (Fig. 6a-b and Extended Data Fig. 9a-c). We hypothesis that the broadness of H3K27ac peaks could suggest two distinct subtypes of SSTEs (Fig. 6a-b and Extended Data Fig. 9a-c). We found that SSTEs with broadened H3K27ac peaks generally exhibit higher eRNA signals compared to those with H3K27ac peaks of comparable broadness (Fig. 6c and Extended Data Fig. 9d-g). The nuclear exosome targeting (NEXT) complex^42,43^, essential for degrading non-coding nuclear RNA, plays a significant role in this context^44^. We discovered that Zcchc8 and Rbm7, two core components of the NEXT complex^43^, are enriched only in SSTEs with H3K27ac peaks of comparable length (Fig. 6b, d-e).

**Fig. 6.**
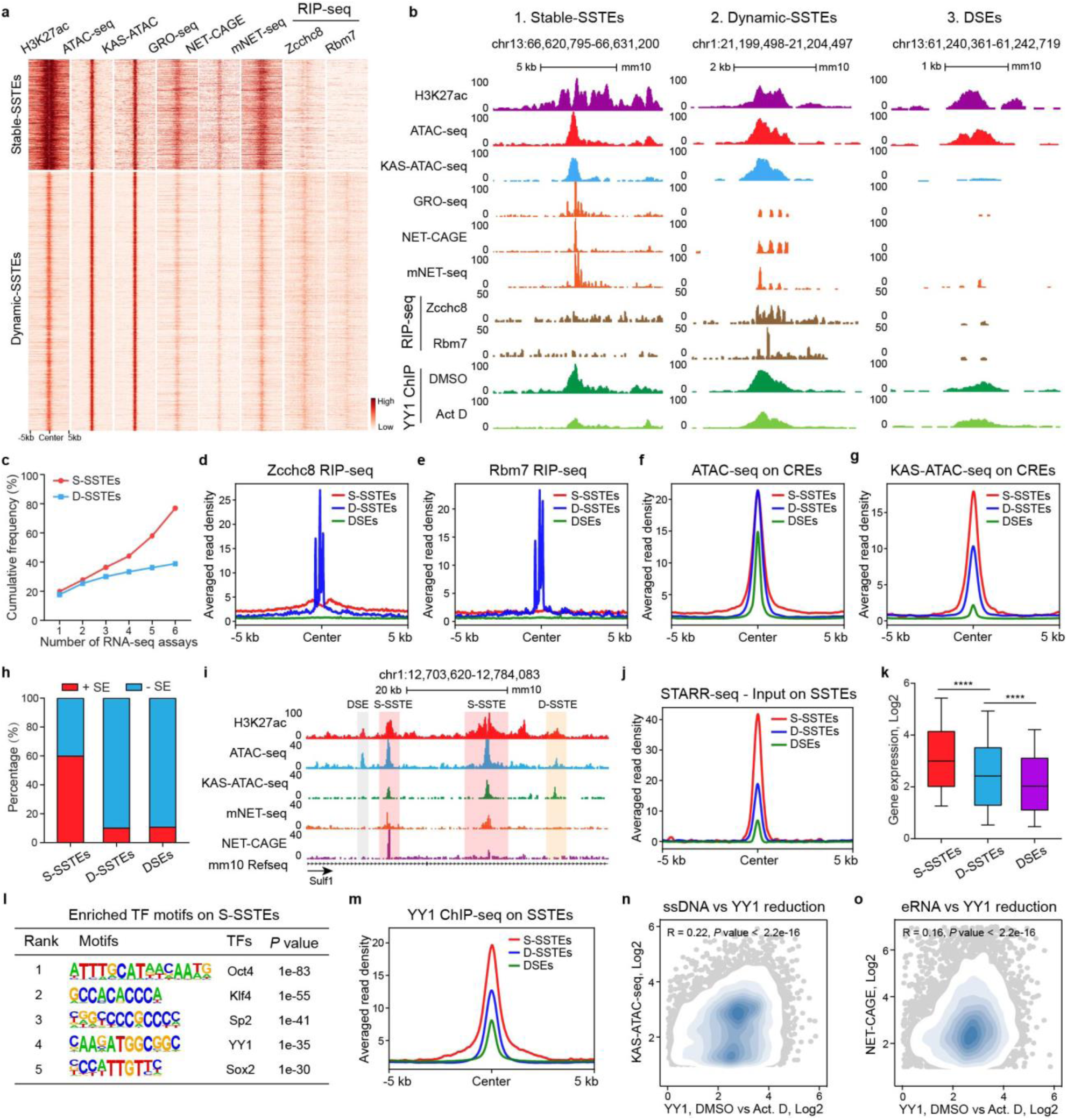
Characteristics and functional implications of two SSTEs types: stable-SSTEs and dynamic-SSTEs. **a**, Heatmap showing the enrichment of H3K27ac ChIP-seq, ATAC-seq, KAS-ATAC-seq, GRO-seq, NET-CAGE, mNET-seq, Zcchc8 RIP-seq, and Rbm7 RIP-seq data on intergenic stable-SSTEs (S-SSTEs, n= 1,961, upper panel) and dynamic-SSTEs (D-SST, n= 11,013, lower panel) in mESCs. S-SSTEs are defined as distal KAS-ATAC-seq peaks spanning less than half the length of H3K27ac peaks, and D-SSTEs are defined as distal KAS-ATAC-seq peaks exceeding half the length of H3K27ac peaks. Regions spanning 5 kb upstream and 5 kb downstream from the center of SSTEs are shown. **b**, Snapshot of UCSC genome browser tracks displaying H3K27ac ChIP-seq, ATAC-seq, KAS-ATAC-seq, GRO-seq, NET-CAGE, mNET-seq, Zcchc8 RIP-seq, Rbm7 RIP-seq, and YY1 ChIP-seq on S-SSTEs (left panel), D-SSTEs (middle panel), and DSEs (right panel). DSEs are identified by distal ATAC-seq peaks without overlapping with KAS-ATAC-seq peaks. YY1 ChIP-seq data from DMSO and Actinomycin D (Act D) treated mESCs are displayed. **c**, Line graph depicting the cumulative frequency of intergenic S-SSTEs (n= 1,961) and D-SSTEs (n= 11,013) with detectable nascent RNA transcripts (FPKM >=0.5) detected by nascent RNA-seq assays, including GRO-seq, PRO-seq, PRO-cap, NET-CAGE, mNET-seq, and 4sU RNA-seq. The red dotted line represents S-SSTEs, and the green dotted line represents D-SSTEs. **d**,**e**, Metagene profiles showing the averaged read density of Zcchc8 RIP-seq (**d**) and Rbm7 RIP-seq (**e**) data across S-SSTEs, D-SSTEs, and DSEs in mESCs. **f**,**g**, Metagene profiles showing the averaged read density of ATAC-seq (**f**) and KAS-ATAC-seq (**g**) data across S-SSTEs, D-SSTEs, and DSEs in mESCs. **h**, Stacked bar plot showing the percentages of S-SSTEs, D-SSTEs, and DSEs that overlap with and without super-enhancers (SEs). **i**, Snapshot of UCSC genome browser tracks displaying H3K27ac ChIP-seq, ATAC-seq, KAS-ATAC-seq, mNET-seq, and NET-CAGE data in mESCs on a representative super enhancer. S-SSTEs within the super enhancer are highlighted in pink, D-SSTEs in orange, and DSEs in grey. **j**, Metagene profile showing STARR-seq signals, with input subtraction, across S-SSTEs, D-SSTEs, and DSEs in mESCs. **k**, Boxplot comparing the transcriptional levels of genes associated with randomly selected 3,000 S-SSTEs, D-SSTEs, and DSEs. The *p* values were calculated using a paired Student’s t-test. The box shows 1st quartile, median and 3rd quartile, respectively. **l**, Table presenting the enriched transcription factors (TFs) motifs identified on S-SSTEs in mESCs. **m**, Metagene profiles showing the averaged read density of YY1 ChIP-seq data across S-SSTEs, D-SSTEs, and DSEs in mESCs. **n**,**o**, Scatterplots showing the Pearson correlation between KAS-ATAC-seq signals (**n**), NET-CAGE signals (**o**), and YY1 binding density reduction caused by Actinomycin D (Act D) treatment on intergenic SSTEs (n=12,601) in mESCs. The Pearson correlation coefficients (R) and associated *p* values are displayed at the top of the scatterplots. Points are color-coded in light blue to indicate peak density.

These findings suggest that eRNA transcripts from SSTEs with extended H3K27ac peaks tend to be more stable, whereas eRNA transcripts from SSTEs with H3K27ac peaks of comparable length appear to be more dynamic and are prone to be degraded by the NEXT complex. We have thus termed SSTEs with extended H3K27ac peaks as stable-SSTEs (S-SSTEs, see Methods) and those with comparable H3K27ac peaks as dynamic-SSTEs (D-SSTEs, see Methods). S-SSTEs are defined as KAS-ATAC-seq peaks that cover over half the length of H3K27ac peaks at distal CREs (Fig. 6a-b and Extended Data Fig. 9a-c), whereas D-SSTEs cover less than half of these H3K27ac peaks (Fig. 6a-b and Extended Data Fig. 9a-c).

In mESCs, we identified 3,247 S-SSTEs, 15,536 D-SSTEs, and 12,999 DSEs (Supplementary Table 2). A higher proportion of S-SSTEs (76.95%, 2,499/3,247) exhibit pronounced eRNA signals compared to D-SSTEs (38.85%, 6,036/15,536) (Fig. 6c). Interestingly, ATAC-seq signals are similar in both subtypes (Fig. 6f). However, S-SSTEs exhibit higher KAS-ATAC-seq read densities and a greater degree of DOI values compared to D-SSTEs (Fig. 6g and Extended Data Fig. 9h). Notably, a substantial proportion of S-SSTEs (60.0%, 1,947/3,247) overlap with super enhancers (SEs) (Fig. 6h-i and Extended Data Fig. 9i), indicating their stronger capacity for gene activation. S-SSTEs also exhibit stronger STARR-seq signals and are associated with higher gene expression levels than D-SSTEs and DSEs (Fig. 6j-k). Overall, our study delineates a novel classification of SSTEs into S-SSTEs and D-SSTEs, expanding our understanding of their molecular characteristics and functional implications in regulating gene expression. Our consensus sequence motifs analysis identified a specific set of TFs enriched on S-SSTEs, including Oct4, Sox2, Nanog, YY1, and c-Myc (Fig. 6l), which are known to play crucial roles in maintaining the self-renewal and pluripotency of mESCs^45,46^. Further analysis of ChIP-seq data for these TFs reveals markedly stronger binding signals on S-SSTEs compared to D-SSTEs and DSEs (Extended Data Fig. 9l), confirming the motif analysis results. In contrast, D-SSTEs displayed enrichment for a distinct set of TFs (Extended Data Fig. 9j). YY1, in particular, exhibits differential binding affinities on S-SSTEs, D-SSTEs, and DSEs (Fig. 6m). Following transcription inhibition using Actinomycin D in mESCs^47^, we found that YY1 binding on S-SSTEs was noticeably reduced but less affected on D-SSTEs and DSEs (Extended Data Fig. 10a-c). This decrease in YY1 binding is significantly correlated with KAS-ATAC-seq and eRNA signals on SSTEs (Fig. 6n-o), indicating that eRNA transcripts and ssDNA play important roles in enhancing the chromatin binding affinity of TFs, particular YY1, on S-SSTEs.

### Difference in topological connectivity preference between SSTEs and DSEs

Topologically Associating Domains (TADs) are fundamental in the higher-order organization of the genome and play crucial roles in transcription regulation^48^. In mESCs, we noticed differences in enrichment of CTCF binding motifs between SSTEs and DSEs. Specifically, CTCF is notably abundant in DSEs, which differs from the TF pattern observed in S- and D-SSTEs (Fig. 7a). In our analysis of ENCODE-defined candidate CREs, including DNase I digestion sites, H3K4me3, distal enhancers, proximal enhancers, promoters, and CTCF binding sites, we intriguingly discovered that a significant proportion (38.7%, 5,032/12,999) of DSEs overlaps with CTCF binding sites (Fig. 7b). Our exploration of the spatial distribution of three CREs subtypes revealed significant differences in topological enrichment preference between S-SSTEs and DSEs. Notably, S-SSTEs are predominantly localized within TADs, while DSEs display a strong localization towards TAD boundaries (Fig. 7c). This distributional preference for DSEs is further shown by their significant enrichment of CTCF and Cohesin binding sequences, proteins indicative of insulators and TAD boundary demarcation (Fig. 7d-f and Extended Data Fig. 11a-c). While previous studies have suggested that transcriptional signals and transcription start sites (TSS) are enriched around topological boundaries^49^, our analysis revealed that CREs within TADs show significantly higher signals in KAS-ATAC-seq and GRO-seq compared to those located at TAD boundaries (Fig. 7e and Extended Data Fig. 11d). Interestingly, CREs located within TADs and at TAD boundaries exhibited comparable levels of ATAC-seq signals (Extended Data Fig. 11e).

**Fig. 7.**
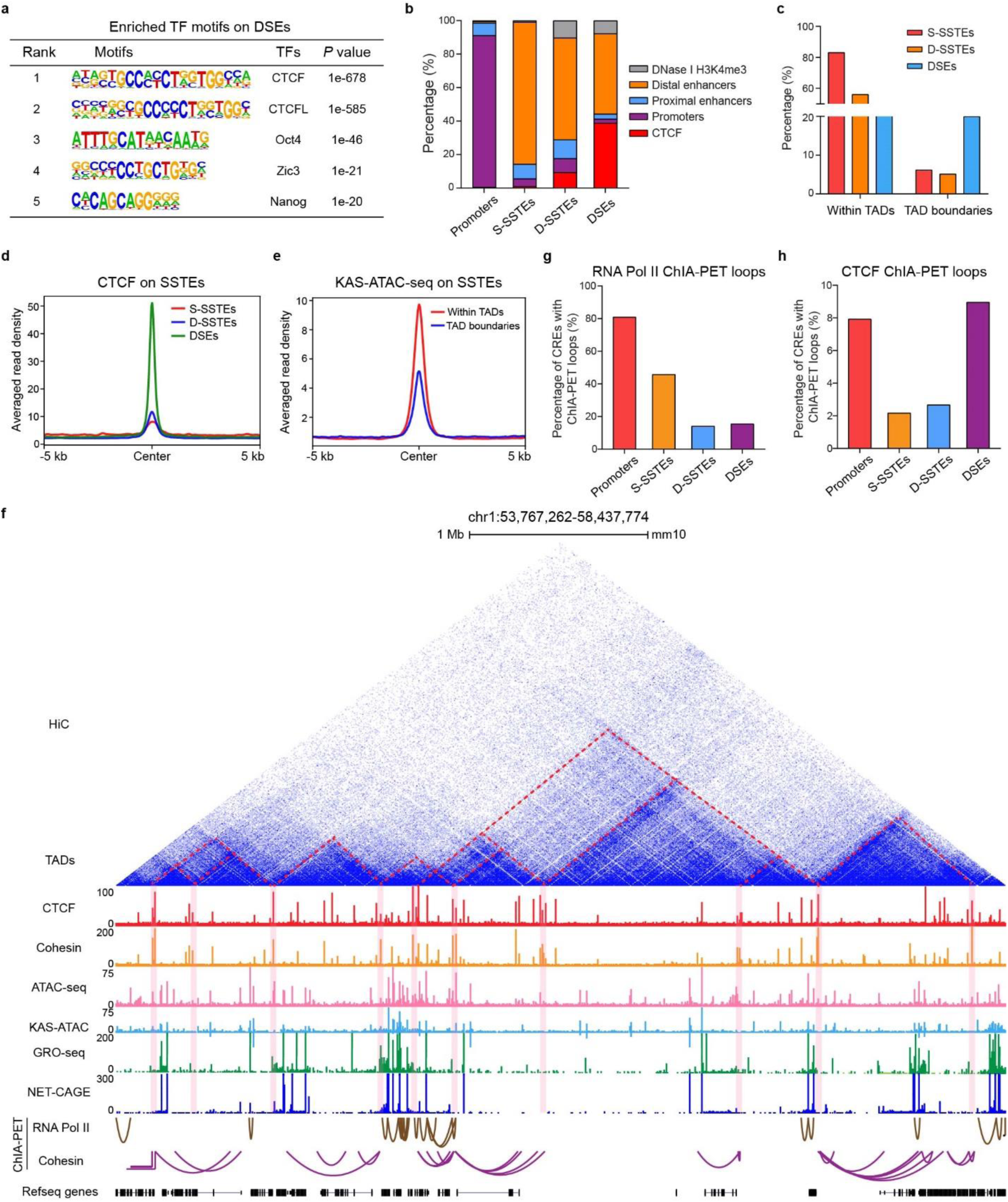
SSTEs and DSEs exhibit distinct distribution preference within higher-order chromatin structures. **a**, Table presenting the enriched transcription factors (TFs) motifs identified on DSEs in mESCs. **b**, Stacked bar plot showing the distribution of different ENCODE-defined candidate cis-regulatory elements (CREs) across M-SSTEs, W-SSTEs, and DSEs in mESCs. The ENCODE-defined candidate CREs include DNase I H3K4me3, distal enhancers, proximal enhancers, promoters, and CTCF binding sites. **c**, Vertical bar plot showing the percentage of S-SSTEs, D-SSTEs, and DSEs located within TADs or at TAD boundaries. **d**, Metagene profiles showing the averaged read density of CTCF ChIP-seq data across S-SSTEs (n=3,247), D-SSTEs (n=15,536), and DSEs (n=12,999) in mESCs. **e**, Metagene profiles showing the averaged read density of KAS-ATAC-seq data across SSTEs within TADs or at TAD boundaries in mESCs. **f**, Snapshot of UCSC genome browser tracks showing the HiC-seq, CTCF ChIP-seq, Cohesin ChIP-seq, ATAC-seq, KAS-ATAC-seq, GRO-seq, NET-CAGE, RNA Pol II and Cohesin ChIA-PET data in mESCs across a representative region (chr1:53,767,262-58,437,774) with prominent topologically associating domains (TADs). TADs are marked by dashed red lines and their boundaries are highlighted in pink. **g**,**h**, Vertical bar plot illustrating the percentages of promoters, S-SSTEs, D-SSTEs, and DSEs linked to long-range interaction loops defined by RNA Pol II (**g**) and CTCF (**h**) ChIA-PET data in mESCs.

To further elucidate the differences in topological connectivity preference between SSTEs and DSEs, we analyzed their intersections with long-range chromatin interaction loops, as defined by RNA Pol II and CTCF ChIA-PET data^7,50^. Our findings indicated that S-SSTEs and promoters are closely associated with RNA Pol II- and YY1-mediated long-range interactions (Fig. 7g and Extended Data Fig. 11f), whereas DSEs primarily align with CTCF- and Cohesin-mediated long-range interactions typically found in insulator regions that delineate TAD boundaries (Fig. 7h and Extended Data Fig. 11g). Taken together, our findings elucidate different spatial distribution of CREs throughout the genome. This distinction is marked by S-SSTEs being more prevalent in intra-TAD regions, while DSEs are inclined towards TAD boundaries. These observations suggest a distinct role in genome organization and gene regulation for S-SSTEs.

## Discussion

In this study, we introduce KAS-ATAC-seq, a new method that combines *Opti-*KAS-seq with ATAC-seq. This approach provides a refined perspective on the dynamic transcriptional landscape of CREs and establishes connections between transcriptional change and function. Our method represents a significant advancement in genomic research as it overcomes limitations of existing methods in terms of accuracy, sensitivity, or input material requirements. It enables a more comprehensive exploration of the regulatory genome.

KAS-ATAC-seq stands out for its dual capability to simultaneously uncover chromatin accessibility and transcriptional activity of CREs. It enables the evaluation of dsDNA openness across ATAC-seq peaks through newly devised DNA Openness Index (DOI). Importantly, KAS-ATAC-seq and the DOI have proven to be more accurate indicators of gene expression compared to ATAC-seq and active histone marks. Furthermore, KAS-ATAC-seq facilitates the de novo annotation of Single-Stranded Transcribing Enhancers (SSTEs) as a subset of distal CREs delineated by ATAC-seq peaks. Our findings reveal that SSTEs are highly enriched with nascent RNA transcription and specific transcription factors (TFs) binding sites, playing a pivotal role in defining stem cell identity.

KAS-ATAC-seq identifies more SSTEs and exhibits greater sensitivity in defining transcribed CREs compared to nascent RNA-based assays. This is partly due to the fact that the eRNAs on transcribed CREs are relatively unstable and can be degraded by the NEXT complex. Notably,

KAS-ATAC-seq also offers a substantial advantage in characterizing transcriptionally active and functional enhancers over ATAC-seq and active histone marks, which tends to identify a considerable number of CREs as insulators or other non-transcribed CREs lacking enhancer activity. This distinction is underscored by the marked enrichment of CTCF and Cohesin binding on DSEs as opposed to SSTEs. Furthermore, SSTEs may offer enhanced sensitivity in predicting functional promoter-enhancer interaction loops compared to ATAC-seq and ChIP-seq peaks of active histone marks (H3K27ac and H3K4me1). Applying KAS-ATAC-seq to the neural differentiation of mESCs into NPCs has revealed intricate details of the transcriptional dynamics of CREs involved in this process. We discovered that KAS-ATAC-seq is more capable of detecting immediate-early transcriptional changes of CREs compared to conventional ATAC-seq, highlighting its advantage in dissecting stem cell neural differentiation and investigating developmental processes. The enhanced resolution offered by KAS-ATAC-seq is likely attributable to its dual capability to simultaneously reveal chromatin accessibility and transcriptional activity of CREs. This also aligns with findings from published studies, which indicate that integrating RNA-seq with ATAC-seq resolves differences better than ATAC-seq alone^51^.

KAS-ATAC-seq also has limitations. For example, the resolution of KAS-ATAC-seq is inherently linked to the quality of chromatin accessibility and ssDNA labeling, which might be influenced by the chromatin state and cellular context. Potential biases introduced during these processes could affect the accuracy and reproducibility of the results. It is also crucial to consider that while KAS-ATAC-seq offers enhanced sensitivity and specificity, it may not capture all aspects of chromatin dynamics, and thus should ideally be used in conjunction with other methods to provide a comprehensive view.

Despite these potential challenges, the ability of KAS-ATAC-seq to provide a detailed and quantitative view of transcriptional regulation opens up new possibilities for understanding complex biological processes.

## Method

### Cell culture

HEK293T cells were purchased from ATCC (CRL11268) and were cultured in DMEM (Gibco 11995) supplemented with 10% (v/v) fetal bovine serum (Gibco), 1% penicillin and streptomycin (Gibco) and grown at 37 °C with 5% CO2. Murine embryonic stem (ES) cells were purchased from ATCC (CRL-1821) and were cultured in DMEM (Gibco 11995) supplemented with 10% (v/v) fetal bovine serum (Gibco), 1 mM L-glutamine (Gibco), 0.1 mM β-mercaptoethanol (Gibco), 1% (v/v) nonessential amino acid stock (100x, Gibco), 1% penicillin/streptomycin stock (100x, Gibco), and 1,000 U/mL LIF (Millipore). Cell lines used in this study were examined for mycoplasma contamination test using LookOut Mycoplasma PCR Kit (Sigma, MP0035).

### *Opti-*KAS-seq

N_3_-kethoxal was synthesized according to an established protocol. Cells harvested from the culture dish or Fluorescence-activated cell sorting (FACS) cells were washed with DPBS and subsequently resuspended in 50 µL of ATAC-Resuspension Buffer (RSB) containing 0.1% NP40, 0.1% Tween-20, and 0.01% Digitonin. After gentle pipetting, the suspension was incubated on ice for 3 minutes. The cells were then treated with 1 mL of cold ATAC-RSB containing 0.1% Tween-20 and centrifuged at 500 RCF for 5 minutes at 4°C. Following a wash with 1 mL DPBS, the cells underwent N_3_-kethoxal labeling at 37 °C with continuous agitation. The labeling process involved a 15-minute incubation in 5mM N_3_-kethoxal in PBS. Genomic DNA (gDNA) was subsequently extracted using the PureLink genomic DNA mini kit. Then, 1 µg genomic DNA was suspended in 95 µL DNA elution buffer supplemented with 5 µL 20 mM DBCO-PEG4-biotin (DMSO solution, Sigma, 760749), 25 mM K3BO3, and incubated at 37 °C for 1.5 h while being gently shaken. Next, 5 µL RNase A (Thermo, 12091039) was added into the reaction mixture followed by incubation at 37 °C for 5 min. Biotinylated gDNA was then recovered by DNA Clean & Concentrator-5 kit (Zymo, D4013). gDNA was suspended into 100 µL water and was fragmented to 150–350 bp size by using Bioruptor Pico at 30s-on/30s-off setting for 30 cycles; 5% of the fragmented DNA was saved as input, and the remaining 95% was used to enrich biotin-tagged DNA by incubation with 10 µL pre-washed Dynabeads MyOne Streptavidin C1 (Thermo, 65001) at room temperature for 15 min. The beads were washed, and DNA was eluted by heating the beads in 15 µL H2O at 95 °C for 10 min. Eluted DNA and its corresponding input were used for library construction by using Accel-NGS Methyl-seq DNA library kit (Swift, 30024).

### *Opti-*KAS-seq using mice tissues

Male B6 mice were purchase from the Jackson Laboratory (catalog no. C57BL/6J). All mice were used at 6-12 weeks of age. Mice were housed under pathogen-free conditions per the National Institutes of Health (NIH) Guide for the Care and Use of Laboratory Animals. All animal care and experiments were approved by the University of Chicago Institutional Animal Care and Use Committee (IACUC), and are compliant with all relevant ethical regulations regarding animal research. Homogenize mouse heart, lung, and spleen tissue to a cell suspension in ice-cold PBS by using a dounce homogenizer or a pellet pestle. Spin the cell suspension at 100 g for 15 seconds to sediment and remove potential large tissue pieces at the bottom of the tube. Spin the cell suspension at 800 g for 5 min. Remove the supernatant and save the cell pellet at the bottom of the tube for labeling. Suspend 5 million cells in 50 µL of ATAC-Resuspension Buffer (RSB) containing 0.1% NP40, 0.1% Tween-20, and 0.01% Digitonin. After gentle pipetting, the suspension was incubated on ice for 3 minutes. The cells were then treated with 1 mL of cold ATAC-RSB containing 0.1% Tween-20 and centrifuged at 500 RCF for 5 minutes at 4°C. Following a wash with 1 mL DPBS, the cells underwent N_3_-kethoxal labeling at 37 °C with continuous agitation. The labeling process involved a 15-minute incubation in 5mM N_3_-kethoxal in PBS. Post-labeling, the cells were treated with a solution of 125 mM H3BO3 (pH 6) and then centrifuged at 500 RCF at room temperature. Isolate total DNA from cells by using PureLink genomic DNA mini kit (Thermo K182001). Elute DNA by using 50 µL 25 mM K3BO3 (pH 7.0). Perform biotinylation and purification, enrichment of N_3_-kethoxal-modified DNA, library preparation and sequencing according to the protocol for mammalian live cells.

### KAS-ATAC-seq

To execute the KAS-ATAC-seq protocol, start by preparing the ATAC-Resuspension Buffer (RSB) by combining 500 µl of 1M Tris-HCl (pH 7.4), 100 µl of 5M NaCl, 150 µl of 1M MgCl2, and then add sterile water up to 50 ml. For the Transposition Mix designated for each 50 µl sample, mix 25 µl of 2x TD buffer, 16.5 µl of PBS, 5 µl of water, 1 µl of 1% (wt/vol) digitonin (added twice for a total of two times), and 2.5 µl of Illumina Tn5 transposase. Initiate the procedure by collecting 50,000 viable cells, wash the collected cells with DPBS and centrifuge at 500 RCF for 5 minutes at 4°C. Resuspend the cell pellet in 50 µl of ATAC-RSB containing 0.1% NP40, 0.1% Tween-20, and 0.01% Digitonin. Mix thoroughly by pipetting up and down three times and then incubate on ice for 3 minutes. Subsequently, dilute this lysed solution with 1 ml of cold ATAC-RSB that has 0.1% Tween-20 (but devoid of NP40 or digitonin). Mix by inverting the tube five times. Centrifuge this mixture again at 500 RCF for 5 minutes at 4°C and discard the supernatant. Resuspend the pellet with 100 µl of 10 mM N_3_-Kethoxial in PBS and then incubate at 37°C with 500 rpm shaking for 10 minutes. After incubation, wash the nuclei by adding 1 ml of DPBS. Centrifuge the suspension at 500 RCF for 5 minutes at 4°C, and subsequently remove the supernatant. Resuspend the nuclei pellet in 50 µl of the previously prepared Transposition Mix. Achieve a uniform mixture by pipetting up and down six times.

Incubate the mixture in a thermomixer set to 37°C and 1,000 rpm for 30 minutes. Remove the tubes from the thermomixer, and immediately terminate the transposition reaction by adding 250 µL (five volumes) of DNA Binding Buffer from the DNA Clean and Concentrator-5 Kit and mix well by pipetting or inversion. Mix thoroughly and followed by brief centrifugation to collect the contents at the bottom of the tubes. Proceed with purifying the transposed DNA using the kit, according to the manufacturer’s instructions. Elute the purified DNA in 21 µl of elution buffer and store at -20°C for future use. Finally, utilize this eluted gDNA for the Click reaction and perform library preparation and sequencing according to the established ATAC-seq protocol^52^.

### ATAC-seq

The ATAC-seq experiments were conducted following the established ATAC-seq protocol (Omni-ATAC-seq) for chromatin accessibility profiling^52^.

### Induced neural differentiation of mESCs into NPCs in vitro

To initiate neural differentiation of mouse embryonic stem cells (mESCs), start by enzymatically digesting the mESCs with 0.05% trypsin to obtain a suspension of single cells. Neutralize the reaction in ESC culture medium without Leukemia Inhibitory Factor (LIF) and centrifuge to remove the supernatant. Resuspend the cells in basal differentiation medium (DMEM supplemented with 15% (v/v) stem cell qualified fetal bovine serum (Gibco), 1 mM GlutaMAX (Gibco), 0.1 mM β-mercaptoethanol (Gibco), 1% (v/v) nonessential amino acid stock (100x, Gibco), 1% penicillin/streptomycin stock) to a density of 1.5×10^6^ cells/ml and cultured at 37°C for 2 days with 5% CO_2_. On Day2, transfer the embryoid bodies to a new dish using basal differentiation medium, allowing them to settle for 2 minutes before discarding the supernatant. Resuspend the embryoid bodies using basal differentiation medium with 1 µM Retinoic Acid (RA). On Day 4, Day 6 and Day 8, wash the cells with DPBS change the medium using basal differentiation medium containing 1 µM RA. After 8-day culture in these conditions, mESCs was differentiated into the neural progenitor cells (NPCs)^41,53^.

### *Opti-*KAS-seq and KAS-seq data processing

To align *Opti-*KAS-seq and KAS-seq data to the reference genome of interest, we utilized the trim_galore package to trim off low-quality sequence, adaptor sequence and primer sequence from single-end or paired-end raw fastq files^54^. Subsequently, we employed bowtie2 to perform the read alignment of *Opti-*KAS-seq and KAS-seq data^55^. Mapped reads in sam files from the aligners are sorted and converted to bam files using ‘samtools sort’^56^, which are subsequently deduplicated using ‘picard MarkDuplicates’ (pair-end data) or ‘samtools rmdup’ (single-end data)^56^. For single-end *Opti-*KAS-seq or KAS-seq data, mapped reads were extended to 150bp as default, irrespective of the initial sequencing data’s read length. For paired-end KAS-seq data, the KAS-Analyzer incorporates a Python script, facilitating the merging of “properly paired” mapped reads into a singular interval^57^. Peak calling for broad KAS-seq was executed using epic2^58^, which identified broad peaks maintaining a false discovery rate (FDR) of 0.05 and a 1.5-fold change relative to the Input. Comprehensive quality control measures, such as library complexity metrics, inter-replicate correlation analyses, fingerprint plots, saturation analyses, genomic distribution of KAS-seq peaks, enrichment within gene-coding regions, and the Fraction of Reads in Peaks (FRiP) metric, were executed with the KAS-Analyzer package. Deduplicated mapped KAS-seq reads were converted to bedGraph and bigWig using deeptools bamCoverage and UCSC tools^59,60^. Intersection analysis between *Opti*-KAS-seq and KAS-seq peaks were performed using bedtools intersect tools. Overlap analysis between *Opti-*KAS-seq and KAS-seq peaks was executed with the bedtools intersect tool^61^.

### KAS-ATAC-seq and ATAC-seq data processing

All data from KAS-ATAC-seq and ATAC-seq were produced in paired-end mode. The preprocessing, read alignment, and read alignments format conversion procedures for KAS-ATAC-seq and ATAC-seq data mirrored those for *Opti*-KAS-seq and KAS-seq. Peak calling for KAS-ATAC-seq and ATAC-seq was accomplished via MACS2^62^, targeting sharp peaks with a false discovery rate (FDR) threshold of 0.05. Deduplicated KAS-seq reads were then transformed into bedGraph and bigWig formats using the deeptools bamCoverage and UCSC tools. Fragment size comparisons within KAS-ATAC-seq and ATAC-seq libraries were facilitated by the ATACseqQC Bioconductor package^63^. Overlap analysis between KAS-ATAC-seq and ATAC-seq peaks was executed with the bedtools intersect tool. Additionally, the distribution of SSTEs on the ENCODE Registry of candidate CREs in the mouse genome was also assessed with the bedtools intersect tool^61,64^.

### HiC data processing

In this study, we analyzed published Hi-C data (GSM2977176 and GSM2977177) in mESCs to elucidate the 3D chromatin organization, employing the Juicer toolkit for comprehensive data processing^64^. Initially, paired-end Hi-C raw data were trimmed to remove adapters and low-quality sequences using trim_galore package^54^, followed by alignment to the mouse reference genome (mm10) using the BWA-MEM aligner integrated within Juicer^64,65^. Subsequent to alignment, duplicate reads were filtered out, and the data were binned into contact matrices to generate a multi-resolution .hic file. MboI restriction sites was used. We then employed HICCUPS (java - Xmx10g -jar juicer_tools_1.22.01.jar hiccups -r 5000,10000 /content/mESCs_30.hic mESCs_hiccups) and Arrowhead algorithms (java -Xmx20g -jar juicer_tools_1.22.01.jar arrowhead -r 10000 -c 20 /content/mESCs_30.hic mESCs_arrowhead) on google colab, also part of the Juicer toolkit, to call significant chromatin loops and Topologically Associating Domains (TADs), respectively. The mESCs .hic files were used to visualize contact matrices in mESCs on the UCSC genome browser^60^.

### Calculation of DNA Openness Index (DOI)

To quantitatively evaluate the dsDNA openness and the associated transcriptional activity throughout the regulome, we formulated the DNA Openness Index (DOI) metric, which is derived by determining the ratio of KAS-ATAC-seq signals to ATAC-seq signals within promoter and distal ATAC-seq peaks. In instances where ATAC-seq peaks exhibited more KAS-ATAC-seq signals than ATAC-seq signals, the ATAC-seq signals were normalized to match the KAS-ATAC-seq signal levels. Specifically, the KAS-ATAC-seq and ATAC-seq signals within ATAC-seq peaks were defined as the count of uniquely mapped reads that overlapped with ATAC-seq peaks by at least 50%.

The DOI can be mathematically calculated as follows:

Where:

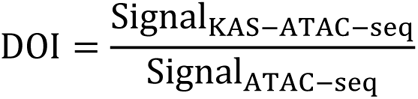

- Signal_KAS-ATAC-seq_ is the quantified read intensity or read count from KAS-ATAC-seq at a specific CRE.
- Signal_ATAC-seq_ is the quantified read intensity or read count from ATAC-seq at the same CRE.

### Identification of Single-Stranded Transcribing Enhancers (SSTEs)

We utilized the KAS-ATAC-seq technique to capture ssDNA generated in accessible chromatin regions. Upon successfully implementing KAS-ATAC-seq in HEK293T and mES cells, we identified KAS-ATAC-seq peaks using MACS2 with default settings. More precisely, we characterized those distal KAS-ATAC-seq peaks that intersected with ATAC-seq peaks as Single-Stranded Transcribing Enhancers (SSTEs). Our findings reveal that SSTEs are distinctively marked by detectable nascent RNA transcription. Notably, we only plot nascent RNA transcription signals on intergenic SSTEs and DSEs to avoid the potential elongation-related transcription signals.

### Classification of SSTEs into stable-SSTEs and dynamic-SSTEs

H3K27ac ChIP-seq peaks exhibited more pronounced length variations compared to KAS-ATAC-seq and ATAC-seq peaks on SSTEs. Based on these observed differences in peak lengths, we categorized SSTEs into two distinct subtypes: stable-SSTEs (S-SSTEs) and dynamic-SSTEs (D-SSTEs). S-SSTEs are defined as SSTEs that cover over 50% of the corresponding H3K27ac peak length on SSTEs. Conversely, D-SSTEs are defined as SSTEs that span less than 50% of the H3K27ac peak length. S-SSTEs typically show significant depletion in RNA transcripts linked to Zcchc8 and Rbm7, whereas D-SSTEs display significant enrichment in RNA transcripts associated with Zcchc8 and Rbm7. Zcchc8 and Rbm7 are integral components of the Nuclear Exosome Targeting (NEXT) complex, which is crucial for the degradation of non-coding nuclear RNA^43,44^. Notably, we only plot nascent RNA transcription signals on intergenic S-SSTEs and D-SSTEs to avoid the potential elongation-related transcription signals.

### Motif analysis for cis-regulatory elements (CREs)

We analyzed consensus sequences and transcription factor binding motifs enriched in various Single-Stranded Transcribing Enhancers (SSTEs) and DSEs using the HOMER findMotifsGenome.pl tool^66^, employing parameters “-len 6,10,13,16 -p 20 -size given -mask”. Figures display p-values corresponding to the ’corrected P’ from the output results.

### Gene Ontology (GO) biological functions using GREAT analysis for SSTEs and DSEs

We employed the Genomic Regions Enrichment of Annotations Tool (GREAT) analysis to predict the potential biological functions associated with SSTEs and DSEs^67^. Bed files containing genomic coordinates of SSTEs and DSEs were uploaded to the GREAT web tool (http://great.stanford.edu/). The tool was set to the default ’Whole Genome’ background with mm10 genome assembly chosen based on the original sequencing data. Association rule settings were kept as default, which assigns genomic regions to nearby genes in a biologically meaningful manner. The output consists of statistically significant annotations from the Gene Ontology (GO) terms were shown.

### Visualization analysis of the TFs binding, histone marks, and nascent RNA transcriptional levels on SSTEs

For the visualized evaluation of the enrichment of transcription factor (TF) binding, histone modifications, and nascent RNA transcription across different subtypes of SSTEs, we employed the deeptools suite to provide a detailed visualization^59^. The plotProfile function was utilized to generate metagene profiles, offering a comprehensive overview of their averaged distribution pattern on SSTEs. In parallel, the plotHeatmap function was engaged to generate heatmap plots, providing an intricate depiction of TF binding intensities, histone marks, and nascent RNA distributions across each specific SSTEs. Furthermore, the UCSC genome browser served as a cloud-based platform, facilitating the visualization of these datasets over selected representative regions. Collectively, these tools provided a profound and cohesive view of the interplay among TF binding, histone modifications, and nascent RNA transcription across diverse SSTEs subtypes.

### Sequence conservation analysis for SSTEs

To investigate the sequence conservation across various SSTEs types in mESC, we employed the evolutionary conservation scores derived from phyloP (phylogenetic p-values) available in the PHAST package (http://compgen.bscb.cornell.edu/phast/). These scores are based on the multiple alignments of 59 vertebrate genomes with the mouse genome. In addition, three alternate sets of scores tailored for specific subsets of species, including Glires, Euarchontoglires, and placental mammals, were also considered. These datasets were sourced from the UCSC genome browser (http://hgdownload.soe.ucsc.edu/goldenPath/mm10/phyloP60way/). Subsequently, to compare the sequence conservation across different SSTEs types in mESC, we employed the plotProfile function within the deeptools suite.

### Statistics and R packages

Scatter plots were made using the ggpubr R package in R (version 3.6.3), which facilitates the creation of beautiful ggplot2-based graphs. The correlation heatmaps that elucidate the relationships among DOI, ATAC-seq, KAS-ATAC-seq, RNA Pol II, and transcription factor binding were constructed using the corrplot package in R (version 3.6.3). Differences in CpG density and transcription factor motif density among SSTEs with varying DOI values were assessed using a two-tailed Student’s t-test. The majority of the bar plots were crafted with GraphPad Prism 7.

### Reporting summary

Further information on research design is available in the Nature Portfolio Reporting Summary linked to this article.

## Code availability

All the bioinformatic scripts used in this study are available at https://github.com/Ruitulyu/KAS-Analyzer. Additionally, the schematic diagrams depicting the *Opti-*KAS-seq and KAS-ATAC-seq experimental procedures were illustrated using the BioRender software.

## Data availability

The raw and processed data of *Opti-*KAS-seq, KAS-seq, KAS-ATAC-seq, and ATAC-seq experiments performed using HEK293T cells and mouse embryonic stem cells have been deposited in the Gene Expression Omnibus (https://www.ncbi.nlm.nih.gov/geo/) under the accession number: GSE256232. All published datasets reanalyzed in this study were summarized in Supplementary Table 3.

## Acknowledgements

We thank the functional genomics facility at the University of Chicago for performing high-throughput sequencing. This work was supported by the US National Institutes of Health (HG006827 to C.H.). C.H. is an investigator of the Howard Hughes Medical Institute. This article is subject to HHMI’s Open Access to Publications policy. HHMI laboratory heads have previously granted a nonexclusive CC BY 4.0 license to the public and a sublicensable license to HHMI in their research articles. Pursuant to those licenses, the author-accepted manuscript of this article can be made freely available under a CC BY 4.0 license immediately upon publication.

## Contribution

C.H. and R.L. conceived the project. R.L. and T.W. developed the *Opti*-KAS-seq and KAS-ATAC-seq experimental procedures and performed KAS-seq, *Opti-*KAS-seq and KAS-ATAC-seq experiments with E14-mESC and HEK293T cells. Y.G. conducted *Opti*-KAS-seq of mouse tissues samples including liver, heart and spleen and the time-course ATAC-seq and KAS-ATAC-seq experiments for the neural differentiation from mouse embryonic stem cells (mESCs) to neural progenitor cells (NPCs) in vitro. P.W. synthesized the N_3_-kethoxal compound to develop *Opti-* KAS-seq and KAS-ATAC-seq. C.Y. performed high-throughput sequencing on KAS-ATAC-seq and ATAC-seq libraries during neural differentiation. R.L. performed the bioinformatic analysis for all the high throughput sequencing data with suggestions from C.Y. R.L. and C.H. wrote the manuscript with input and edits from all authors.

---

**Extended Data Fig. 1.**
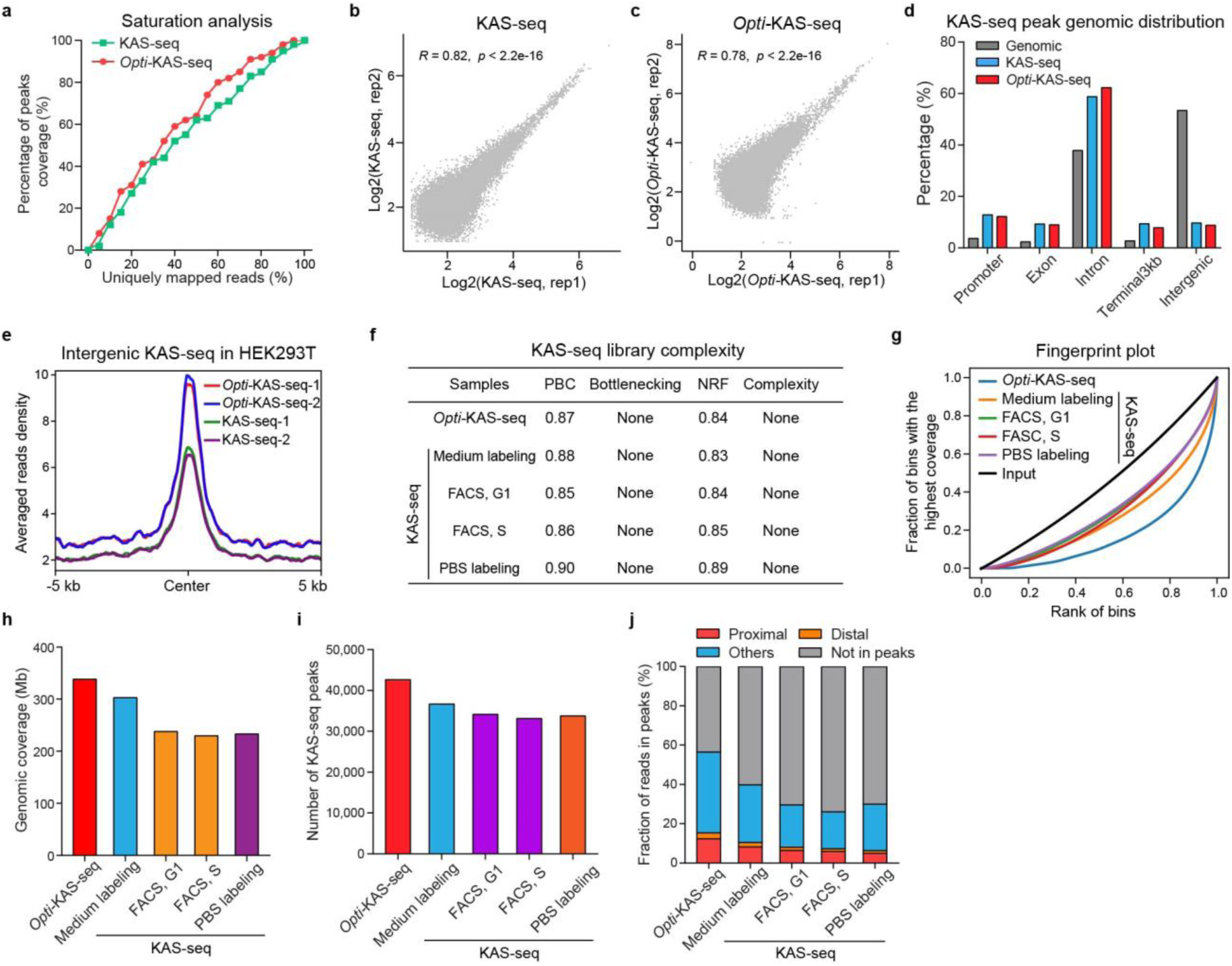
Comparing the quality control of KAS-seq and *Opti*-KAS-seq datasets in HEK293T cells. **a**, Dotted line plot showing the saturation analysis of conventional KAS-seq and Optimized KAS-seq (*Opti-*KAS-seq) data in HEK293T cells. **b**,**c**, Scatterplots showing the Pearson correlation between two replicates of KAS-seq (**b**) and *Opti-*KAS-seq (**c**) data in HEK293T cells. Grey dots represent KAS-seq peaks (**b**, n=54,073; **c**,=72,815). Both the Pearson correlation coefficients and significance (*p* values) are provided. **d**, Grouped bar plot illustrating the distribution of KAS-seq and *Opti-*KAS-seq peaks across various genomic features (promoter, exon, intron, terminator, and intergenic regions) in HEK293T cells. Promoters are defined as regions 2 kb upstream and downstream from the transcription start site (TSS). The relative proportions of each genomic feature, based on the hg19 Refseq annotation, are presented as ’Genomic’. **e**, Metagene profiles showing the averaged read density of *Opti-*KAS-seq and KAS-seq data across intergenic KAS-seq peaks (n=2,446) in HEK293T cells. **f**, Table chart showing the library complexity metrics calculated using *Opti-*KAS-seq and KAS-seq data generated in HEK293T cells. **g**, Fingerprint plot of KAS-seq data from different protocols in HEK293T cells showing that the ssDNA capturing efficiency of *Opti-*KAS-seq is sig increased compared to KAS-seq. **h**, Vertical bar plot showing the genomic coverage of KAS-seq peaks from *Opti-*KAS-seq and KAS-seq data generated in HEK293T cells. **I**, Vertical bar plot showing the number of KAS-seq peaks from *Opti-*KAS-seq and KAS-seq data generated in HEK293T cells. **j**, Stacked bar plot showing the fraction of reads in peaks (fRiP), calculated from KAS-seq and *Opti-*KAS-seq reads uniquely mapped to promoters (±500 bp of TSS), distal cis-regulatory elements (CREs) (>500 bp from TSS) and other regions in HEK293T cells. Each bar represents two merged technical replicates. All values were determined from 30 million randomly selected mapped reads.

**Extended Data Fig. 2.**
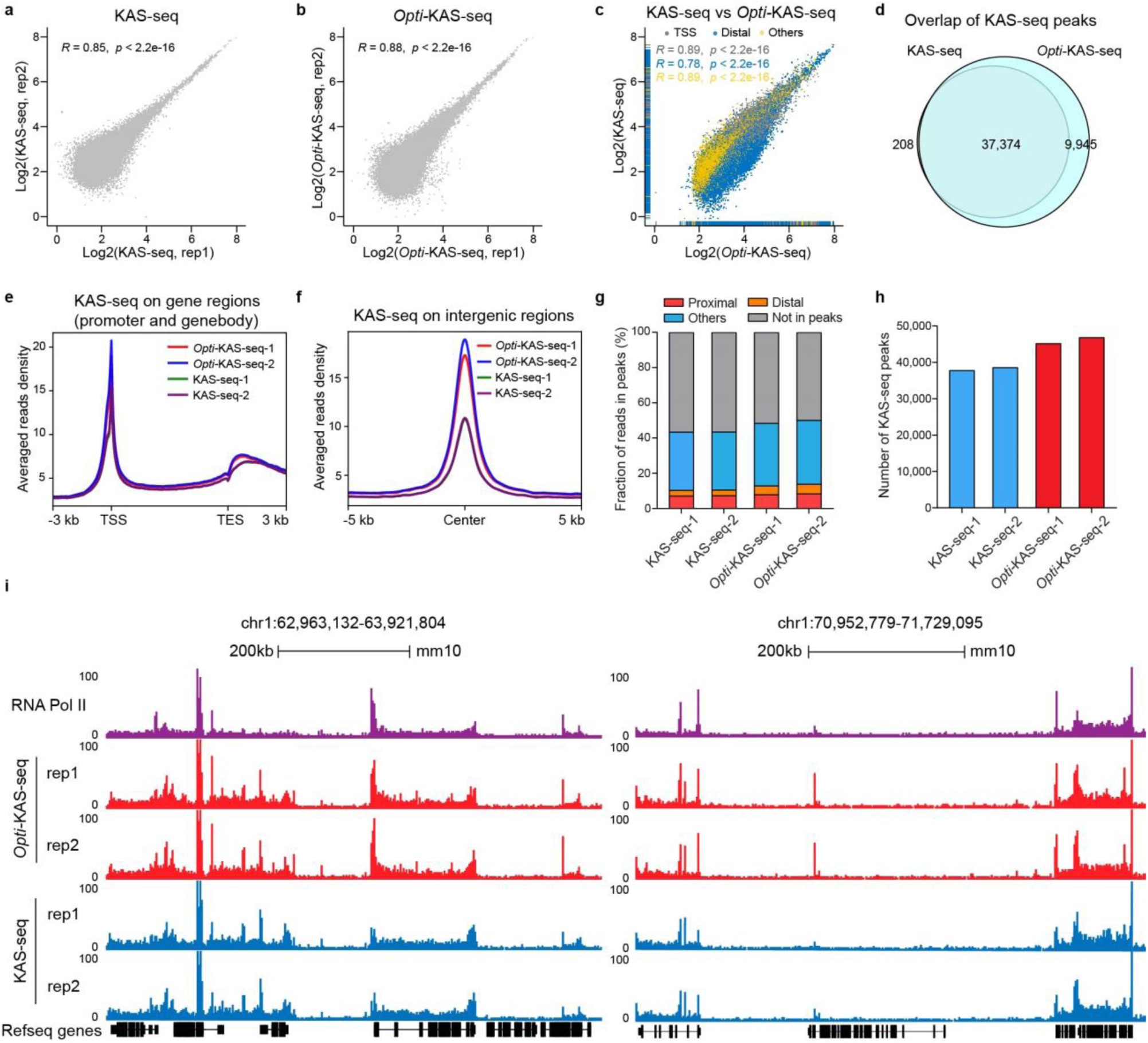
KAS-seq and *Opti-*KAS-seq datasets in mESCs. **a**,**b**, Scatterplot showing the Pearson correlation between two replicates of KAS-seq (**a**) and *Opti-*KAS-seq (**b**) data in mESCs. Grey dots represent KAS-seq peaks (**a**, n= 64,019; **b**, n= 70,037). Both the Pearson correlation coefficients and significance (*p* values) are provided. **c**, Scatterplot comparing KAS-seq and *Opti-*KAS-seq data in mESCs across 1kb genomic bins on merged KAS-seq and *Opti-*KAS-seq peaks. Grey dots denote promoters (n=13,890), blue dots denote distal CREs (n=23,881), and yellow dots denote other genomic bins (n=31,599) (as labeled). Both the correlation coefficients and significance (*p* values) are provided. **d**, Venn diagram illustrating the overlap of peaks identified using KAS-seq and *Opti-*KAS-seq datasets in mESCs. **e**,**f**, Metagene profiles showing the averaged read density of KAS-seq and *Opti-*KAS-seq datasets across gene-coding regions (**e**, n=22,868) and intergenic KAS-seq peaks (**f**, n=12,992) in mESCs. Gene-coding regions were defined as 3 kb upstream of TSS and 3 kb downstream of TES. **g**, Stacked bar plot showing the fraction of reads in peaks (FRiP) that were mapped to promoters (±500 bp of TSS), distal cis-regulatory elements (distal CREs, >500 bp from TSS) and other regions from KAS-seq and *Opti-*KAS-seq datasets generated in mESCs. All values were determined from 30 million random aligned de-duplicated reads. **h**, Vertical bar plot showing the number of KAS-seq peaks from KAS-seq and *Opti-*KAS-seq datasets generated in mESCs. **i**, Snapshot of UCSC genome browser tracks displaying RNA Pol II ChIP-seq, KAS-seq, and *Opti-*KAS-seq data in mESCs across two representative regions. Two replicates for KAS-seq and *Opti-*KAS-seq datasets are displayed.

**Extended Data Fig. 3.**
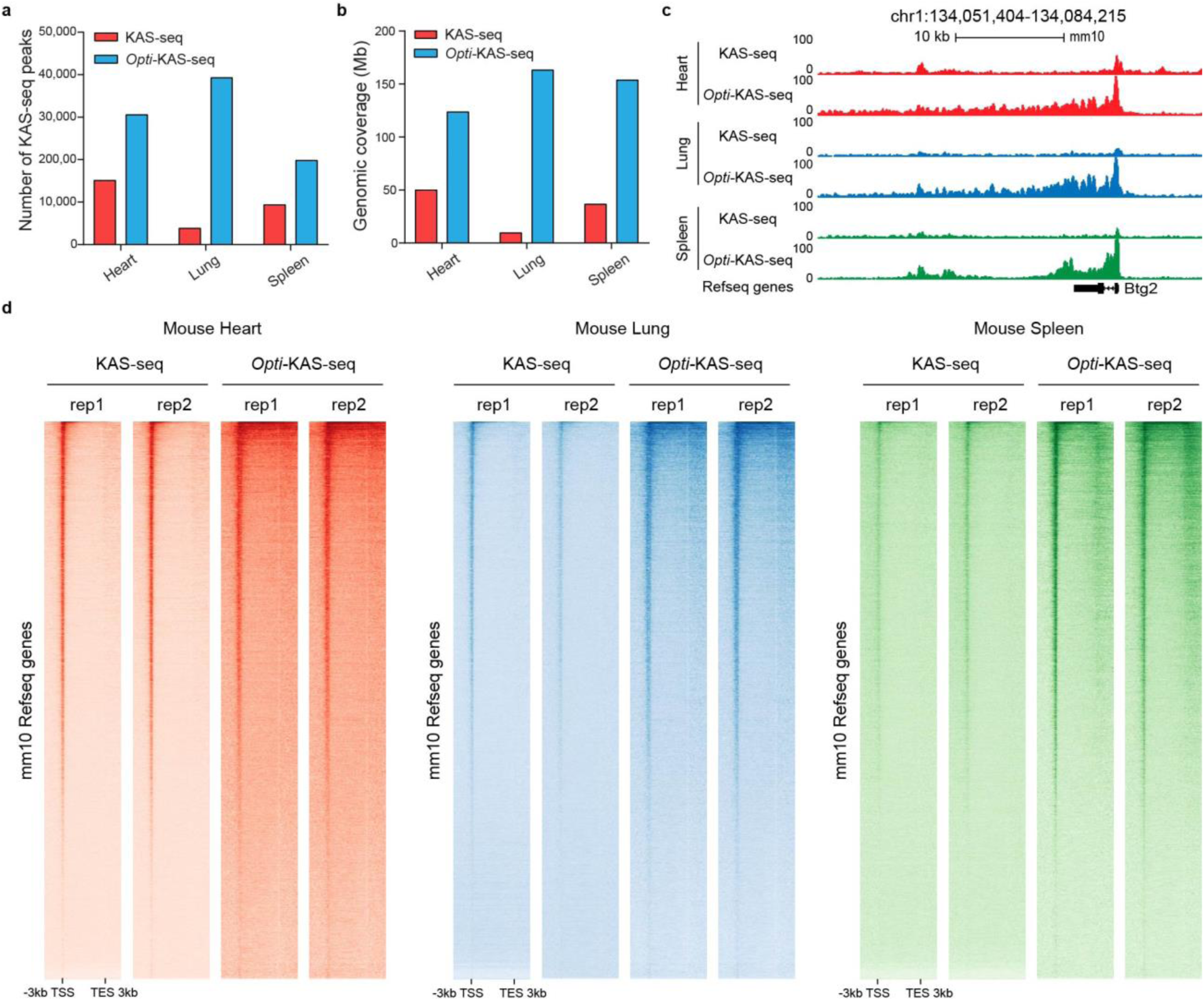
*Opti-*KAS-seq datasets were generated in three mouse tissue types that are not compatible with conventional KAS-seq protocol. **a**, Grouped bar plot displaying the number of KAS-seq peaks from KAS-seq and *Opti-*KAS-seq datasets in mouse heart, lung, and spleen tissues. **b**, Grouped bar plot displaying the genomic coverage of KAS-seq peaks from KAS-seq and *Opti-*KAS-seq datasets generated in mouse heart, lung, and spleen tissues. **c**. Snapshot of UCSC genome browser tracks displaying KAS-seq and *Opti-*KAS-seq datasets generated in mouse heart, lung, and spleen tissues across a representative region (chr1:134,051,404-134,084,215). **d**, Heatmap plots showing the distribution of KAS-seq and *Opti-*KAS-seq read density at gene-coding regions (n=22,868), with 3 kb upstream of TSS and 3 kb downstream of TES shown. KAS-seq and *Opti-*KAS-seq datasets were obtained from mouse heart (left panel), lung (middle panel), and spleen (right panel) tissues.

**Extended Data Fig. 4.**
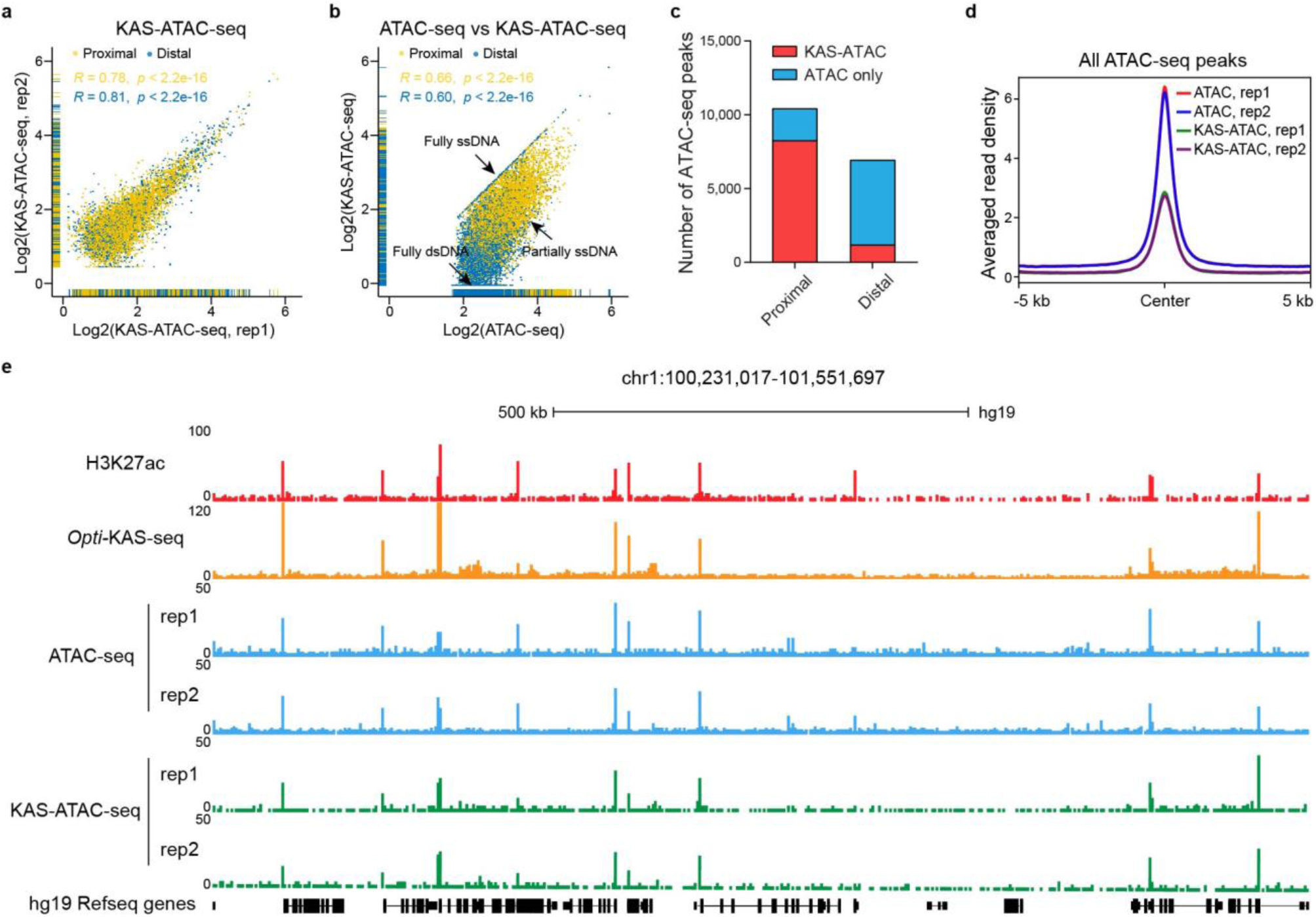
Implementation of KAS-ATAC-seq in HEK293T cells. **a**, Scatterplot illustrating the Pearson correlation between two replicates of KAS-ATAC-seq data for proximal (n=6,863) and distal (n=1,445) peaks in fresh HEK293T cells. Yellow dots represent proximal peaks, blue dots represent distal peaks. Both the Pearson correlation coefficients and significance (*p* values) for proximal and distal peaks are provided. **b**, Scatterplot illustrating the Pearson correlation between ATAC-seq and KAS-ATAC-seq data for proximal (n=9,410) and distal (n=6,913) peaks in fresh HEK293T cells. Yellow dots represent proximal peaks, blue dots represent distal peaks. Both the Pearson correlation coefficients and significance (*p* values) for proximal and distal peaks are provided. Peaks exhibiting higher signals in KAS-ATAC-seq relative to ATAC-seq are normalized to the ATAC-seq signals. **c**, Stacked bar plot showing the numbers of ATAC-seq peaks, depicted by their overlap with KAS-ATAC-seq peaks, in HEK293T cells. Proximal and distal ATAC-seq peaks are shown separately. **d**, Metagene profile showing the distribution of averaged ATAC-seq and KAS-ATAC-seq read density across all ATAC-seq peaks identified in HEK293T cells, with 5 kb upstream and 5 kb downstream from the center of ATAC-seq peaks shown. **e**, Snapshot of UCSC genome browser tracks displaying H3K27ac ChIP-seq, *Opti-*KAS-seq, ATAC-seq, and KAS-ATAC-seq data in HEK293T cells on a representative region (chr1:100,231,017-101,551,697). Two replicates of KAS-seq and *Opti-* KAS-seq datasets are shown.

**Extended Data Fig. 5.**
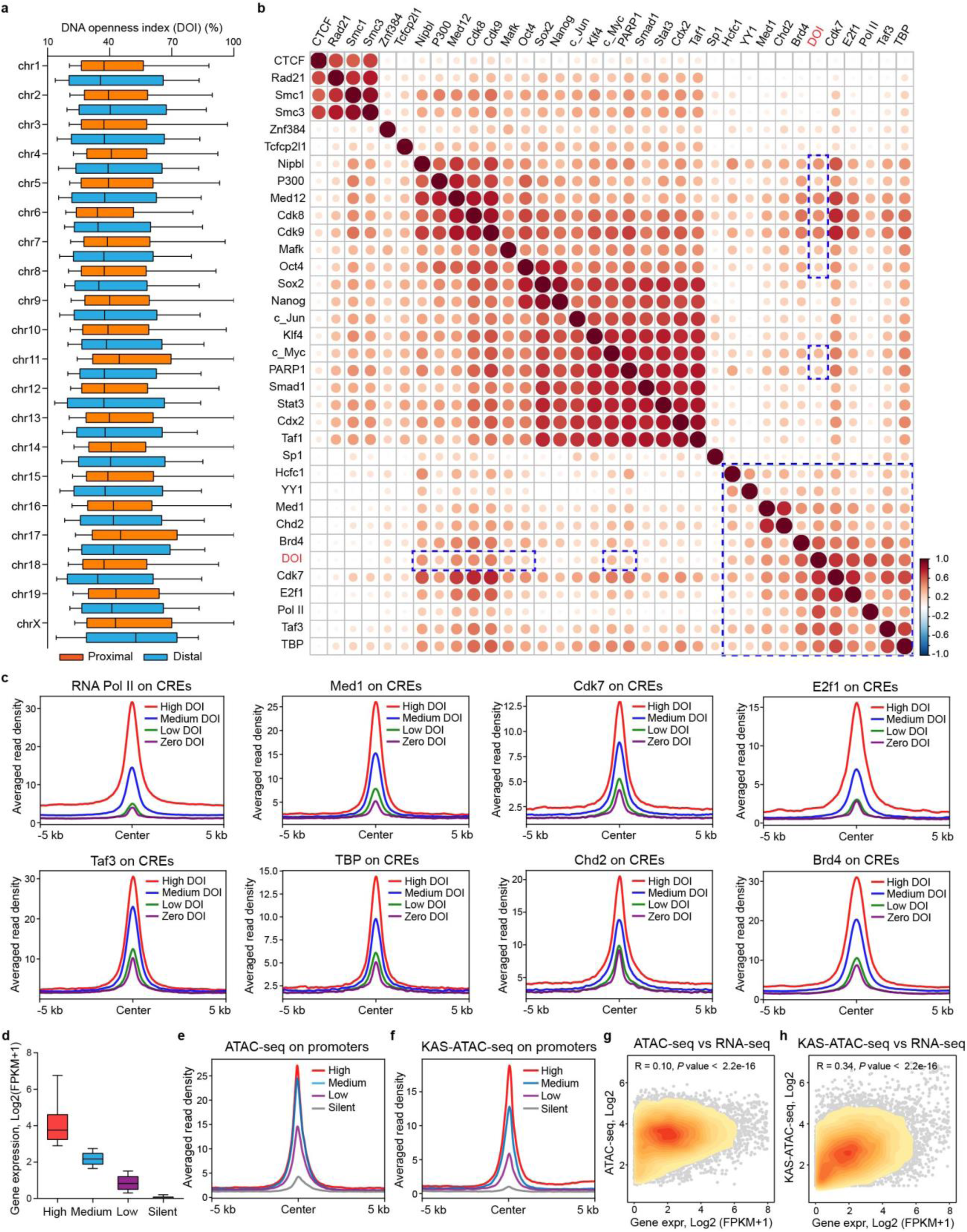
Specific transcription factors (TFs) are highly enriched at cis-regulatory elements (CREs) with high DNA openness index (DOI) in mESCs. **a**, Horizontal boxplot illustrating the DNA openness index (DOI) of proximal and distal cis-regulatory elements (CREs) across different chromosomes. **b**, Heatmap illustrating the Pearson correlation between DNA openness index (DOI), RNA Pol II, and selected transcription factors (TFs) at proximal cis-regulatory elements. TFs that have a Pearson correlation higher than 0.3 with the DNA openness index (DOI) are marked with blue dashed rectangle. **c**, Metagene profiles showing the distribution of RNA Pol II and specific TFs (Med1, Cdk7, E2f1, Taf3, TBP, Chd2, and Brd4) binding density across four categories of proximal CREs in mESCs, with 5 kb upstream and 5 kb downstream from the center of CREs shown. These CREs are categorized based on different levels of DOI (High DOI, Medium DOI, Low DOI, and Zero DOI). **d**, Boxplot illustrating the classification of gene groups based on expression levels (FPKM, calculated using bulk RNA-seq data in mESCs). Genes with FPKM values above 0.5 are classified as expressed genes and then categorized into high, medium, and low expression groups, each group containing an equal number of genes (n=3,822). Genes with FPKM values below 0.5 are classified as silent genes (n=9,047). The box shows 1st quartile, median and 3rd quartile, respectively. **e**,**f**, Metagene profiles showing the distribution of averaged ATAC-seq (**e**) and KAS-ATAC-seq (**f**) read density at gene promoters with different expression levels (high, medium, low, silent) in mESCs, with 5 kb upstream and downstream of TSS shown. **g**,**h**, Scatterplots showing the Pearson correlation between gene expression (Gene expr, bulk RNA-seq) and chromatin accessibility from ATAC-seq (**g**), as well as ssDNA signals from KAS-ATAC-seq (**h**) on expressed genes (n=11,467). The Pearson correlation coefficient (R) and associated *p* values are displayed at the top of the scatterplots. Points are color-coded in orange to indicate gene density.

**Extended Data Fig. 6.**
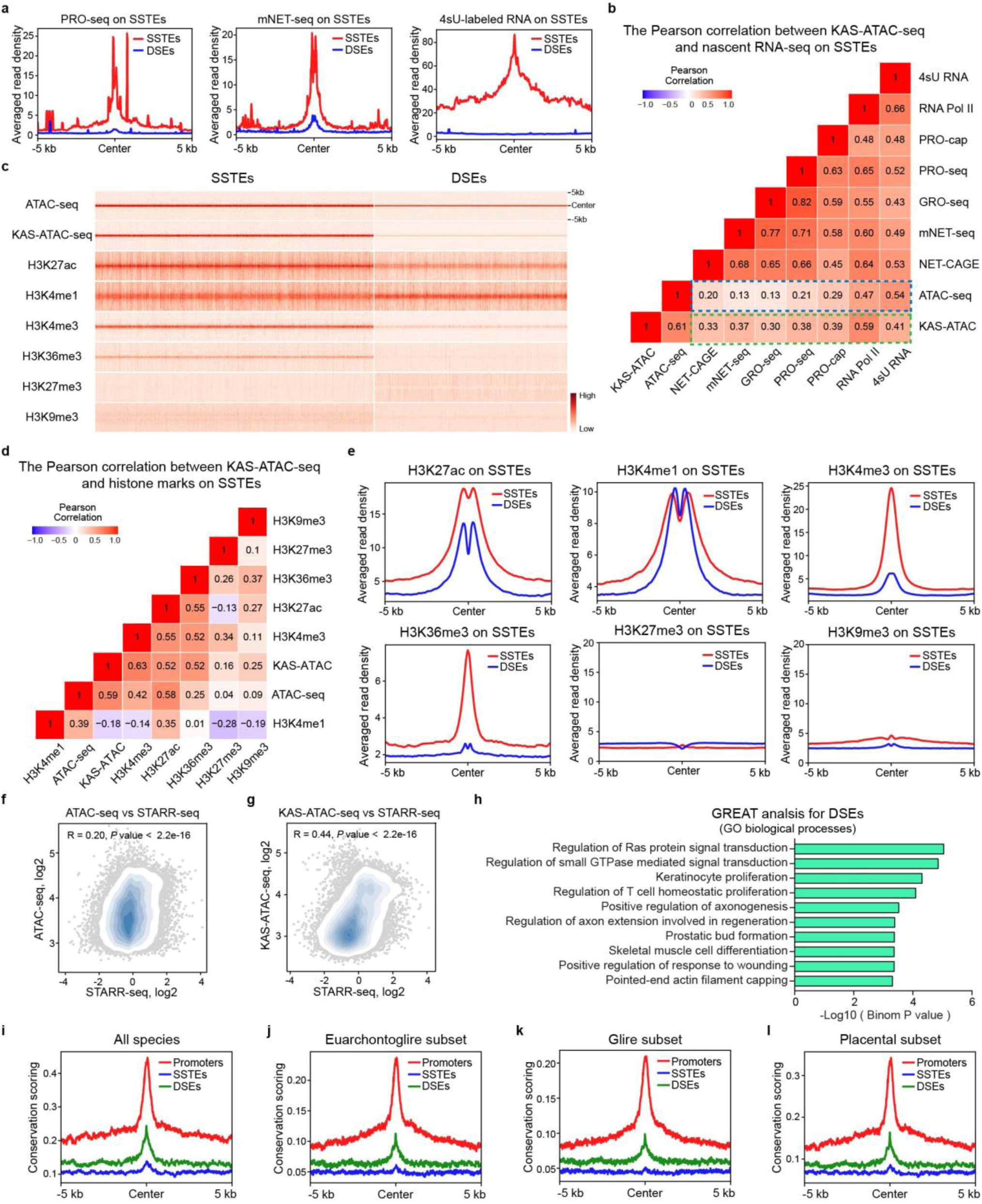
Correlation of KAS-ATAC-seq signals with nascent RNA transcription, histone marks, enhancer activities, and evolutionary conservation score on SSTEs in mESCs. **a**, Metagene profiles showing nascent RNA transcriptional levels determined by various assays, including PRO-seq, mNET-seq, and 4sU RNA-seq, across SSTEs (red) and DSEs (blue), with 5 kb upstream and 5 kb downstream from the center of distal cis-regulatory elements (CREs) shown. **b**, Correlation heatmap illustrating the Pearson correlation coefficients to depict relationships among ATAC-seq, KAS-ATAC-seq, RNA Pol II binding, and nascent RNA transcription (4sU RNA, GRO-seq, PRO-seq, PRO-cap, mNET-seq, NET-CAGE) associated with SSTEs in mESCs. The Pearson correlation coefficients were labeled on the correlation heatmap. Pearson correlation coefficients between ATAC-seq signals and nascent RNA transcription levels are marked with blue dashed rectangles, and while Pearson correlation coefficients between KAS-ATAC-seq signals and RNA transcription levels are marked with green dashed rectangles. **c**, Horizontal heatmap showing the enrichment of ATAC-seq, KAS-ATAC-seq, and various histone marks (H3K27ac, H3K4me1, H3K4me3, H3K36me3, H3K27me3, and H3K9me3) ChIP-seq read density on SSTEs (top panel) and DSEs (bottom panel) in mESCs. Regions spanning 5 kb upstream and 5 kb downstream from the center of distal CREs are shown. **d**, Correlation heatmap illustrating the Pearson correlation coefficients among ATAC-seq, KAS-ATAC-seq, and various histone marks (H3K27ac, H3K4me1, H3K4me3, H3K36me3, H3K27me3, and H3K9me3) ChIP-seq data on SSTEs in mESCs. The Pearson correlation coefficients were labeled on the correlation heatmap. **e**, Metagene profiles showing the averaged read density of various histone marks ChIP-seq data (H3K27ac, H3K4me1, H3K4me3, H3K36me3, H3K27me3, and H3K9me3) over SSTEs (red) and DSEs (blue) in mESCs. **f**,**g**, Scatterplots showing the Pearson correlation between STARR-seq and ATAC-seq (**f**) or KAS-ATAC-seq (**g**) datasets on SSTEs (n=18,783). The Pearson correlation coefficients (R) and associated *p* values are displayed at the top of the scatterplots. Points are color-coded in light blue to indicate SSTE density. **h**, Horizontal bar plot illustrating the Gene Ontology (GO) biological processes derived from the GREAT analysis of DSEs in mESCs. **i**,**j**,**k**,**l**, Metagene profiles showing the averaged read density of evolutionary conservation scores obtained through phyloP methods, for all species (**i**: vertebrates) and three subsets (**j**: Glires; **k**: Euarchontoglires; and **l**: Placental mammal) across promoters (red), SSTEs (blue) and DSEs (green), with 5 kb upstream and 5 kb downstream from the center of CREs shown.

**Extended Data Fig. 7.**
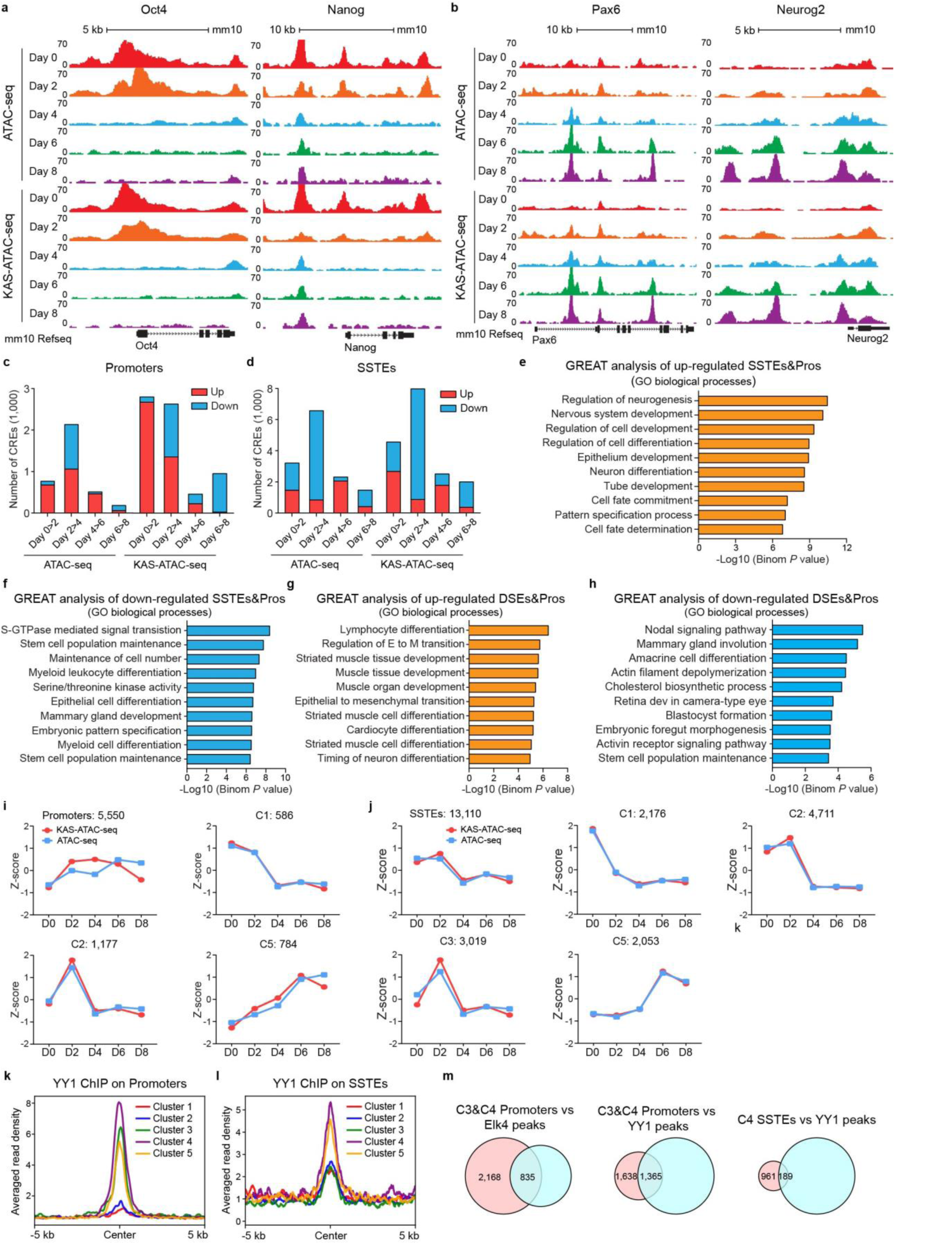
KAS-ATAC-seq accurately captures the transcriptional dynamics of promoters and SSTEs during the neural differentiation of mESCs into NPCs. **a**,**b**, Snapshots of UCSC genome browser tracks showing ATAC-seq and KAS-ATAC-seq data for stem cell pluripotency genes (Oct4 and Nanog) (**a**) and early neural marker genes (Pax6 and Neurog2) (**b**), at specific time point during mouse neural differentiation from mESCs to NPCs. **c**,**d**, Stacked bar plots showing the number of dynamic promoters (**c**) and SSTEs (**d**) identified using ATAC-seq and KAS-ATAC-seq data at different time-points compared to the previous time-point during in vitro neural differentiation from mESCs to NPCs. **e**,**f**, Horizontal bar plot illustrating the Gene Ontology (GO) biological processes derived from the GREAT analysis of up-(**e**) and down-regulated (**f**) ssDNA promoters and SSTEs identified using KAS-ATAC-seq data during mouse neural differentiation from mESCs to NPCs. **g**,**h**, Horizontal bar plot illustrating the Gene Ontology (GO) biological processes derived from the GREAT analysis of up-(**g**) and down-regulated (**h**) non-ssDNA promoters and DSEs identified using ATAC-seq and KAS-ATAC-seq data during mouse neural differentiation from mESCs to NPCs. **i**, Line graph depicting the z-scores of all, C1, C2, and C5 promoters in the clustered heatmap of dynamic promoters. The z-scores were calculated using ATAC-seq data (blue) and KAS-ATAC-seq data (red) at various stages of mouse neural differentiation. **j**, Line graph depicting the z-scores of all, C1, C2, C3, and C5 SSTEs in the clustered heatmap of dynamic SSTEs. The z-scores were calculated using ATAC-seq data (blue) and KAS-ATAC-seq data (red) at various stages of mouse neural differentiation. **k**,**l**, Metagene profiles showing the averaged read density of YY1 ChIP-seq data over different ssDNA promoters (**k**) and SSTEs (**l**) clusters in the clustered heatmaps during mouse neural differentiation. **m**, Venn plots showing the overlap analysis between C3&C4 promoters and Elk4 peaks (left panel), C3&C4 promoters and YY1 peaks (middle panel), and C4 SSTEs and YY1 peaks (right panel).

**Extended Data Fig. 8.**
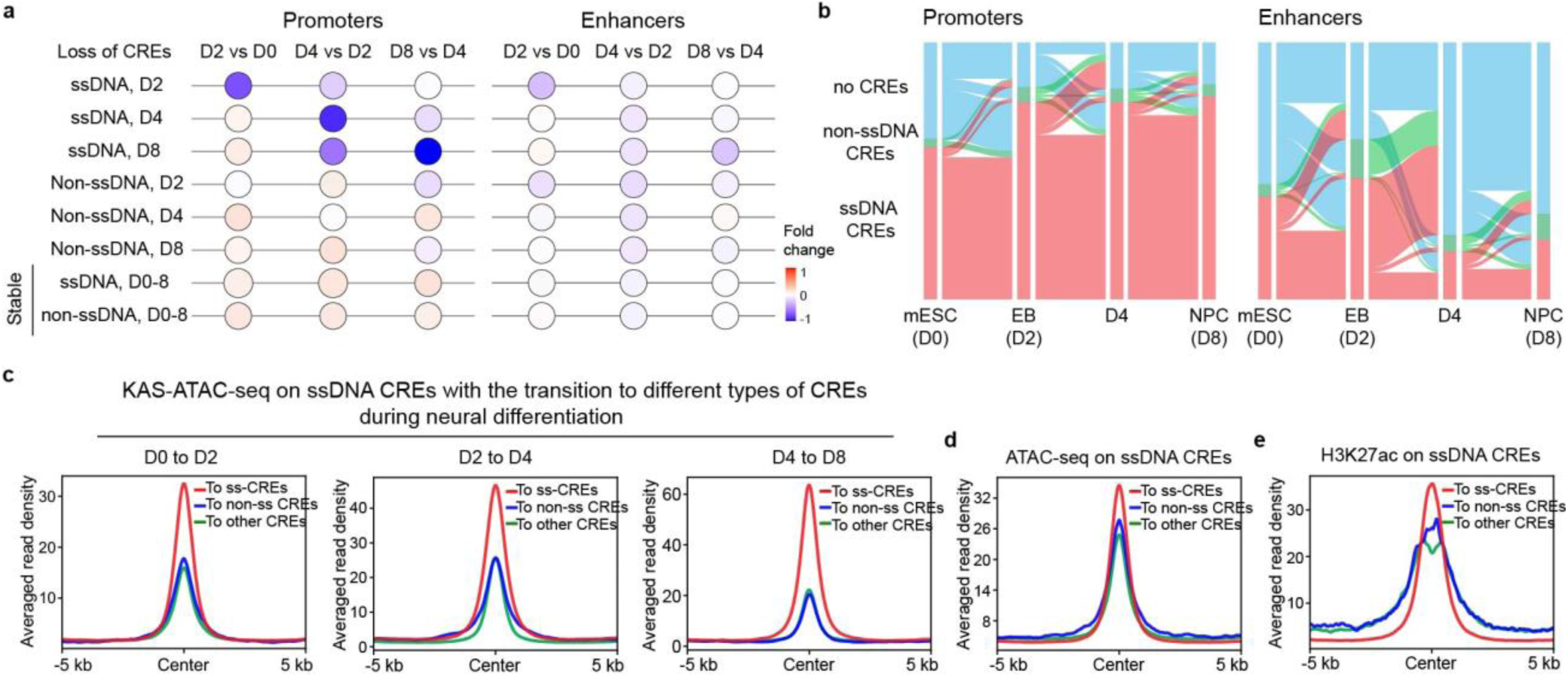
The transition of ssDNA promoters and SSTEs during the neural differentiation of mESCs into NPCs. **a**, Bubble plots showing the fold changes of gene expression levels between consecutive stages for genes associated with “loss of” and stable (no-change) ssDNA and non-ssDNA CREs defined by KAS-ATAC-seq and ATAC-seq data during the neural differentiation from mESCs to EBs and NPCs (D0 to D8). CREs are shown separately as promoters and enhancers. The color key, ranging from blue to red, indicates the median of fold changes of gene expression levels from low to high, respectively. The bulk cell RNA-seq data are referred from GSE151900. **b**, Alluvial plots showing the transition of dynamic promoters (left) and enhancers (right) during the neural differentiation. Each line in the plot represents a dynamic promoter or enhancer, and the total dynamic CREs shown are those classified as dynamic promoter or enhancer in at least one of the analyzed stages. **c**, Metagene profiles showing the averaged read density of KAS-ATAC-seq data over three groups of ssDNA CREs transitioning into various CRE subtypes (to ssDNA CREs, to non-ssDNA CREs, and to other CREs) in later stage of neural differentiation from mESCs to NPCs. **d**,**e**, Metagene profiles showing the averaged read density of ATAC-seq (**d**) and H3K27ac (**e**) ChIP-seq data in mESCs over three groups of SSTEs transitioning into various CRE subtypes (to ssDNA CREs, to non-ssDNA CREs, and to other CREs) during the neural differentiation from mESCs to NPCs.

**Extended Data Fig. 9.**
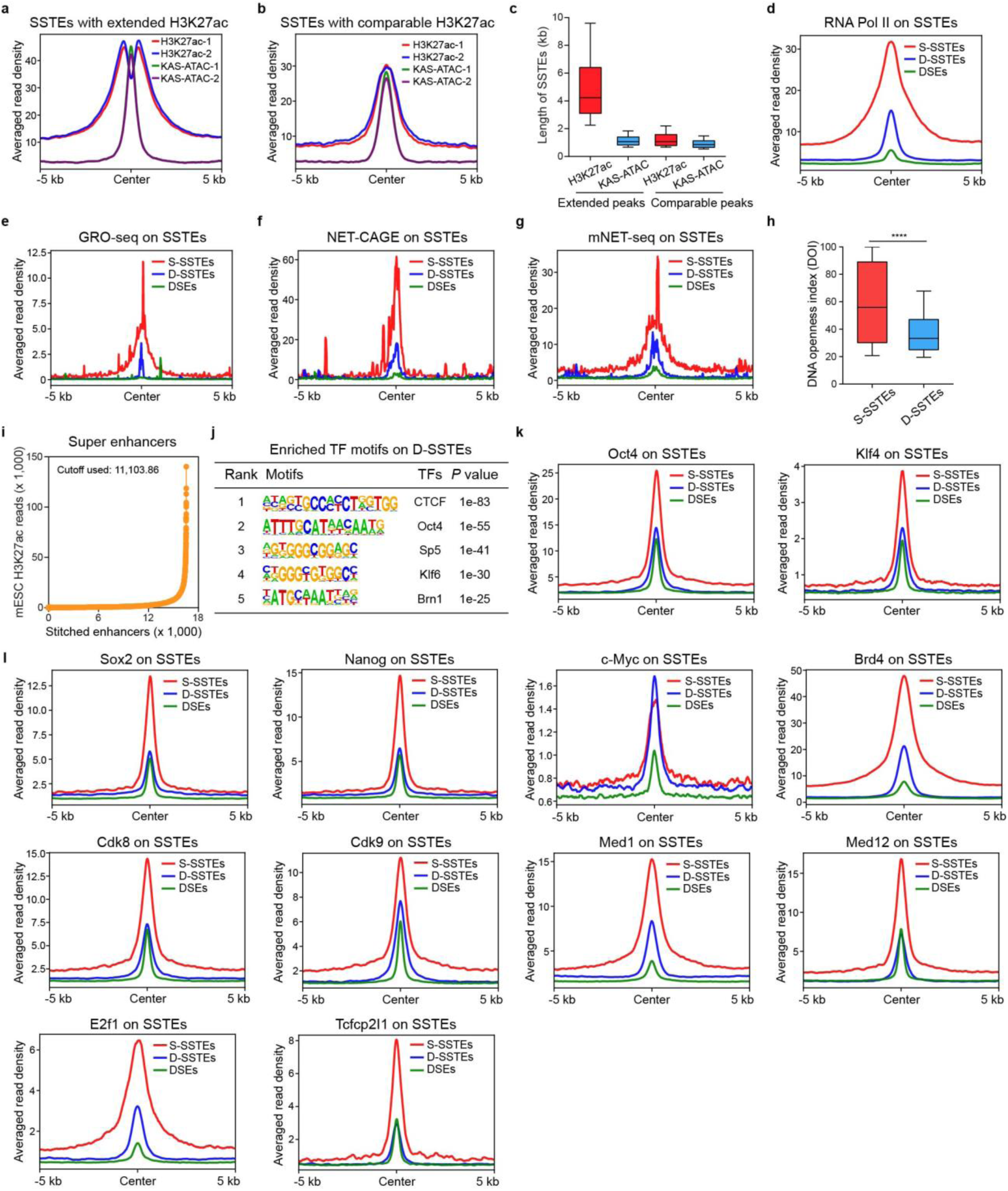
Distinct characteristics and functional implications of two types of SSTE (S- and D-SSTEs) in mESCs. **a**,**b**, Metagene profiles showing the averaged read density of H3K27ac ChIP-seq and KAS-ATAC-seq data across SSTEs with extended H3K27ac peaks (**a**), as well as comparable H3K27ac peaks (**b**) in mESCs. Regions spanning 5 kb upstream and 5 kb downstream from the center of cis-regulatory elements (CREs) are shown. **c**, Box plot comparing the length of H3K27ac, and KAS-ATAC-seq peaks on SSTEs with extended H3K27ac peaks, as well as comparable H3K27ac peaks in mESCs. The box shows 1st quartile, median and 3rd quartile, respectively. **d**,**e**,**f**,**g**, Metagene profiles showing the averaged read density of RNA Pol II (**d**), GRO-seq (**e**), NET-CAGE (**f**), and mNET-seq (**g**) data across S-SSTEs, D-SSTEs, and DSEs in mESCs. Regions spanning 5 kb upstream and 5 kb downstream from the center of SSTEs are shown. **h**, Boxplot comparing the DNA openness index (DOI) of S-SSTEs and D-SSTEs in mESCs. A paired student t-test was used to calculate the p-value. The box shows 1st quartile, median and 3rd quartile, respectively. **i**, Distribution visualization of H3K27ac ChIP-Seq signals identifying 2,646 super enhancers (SEs) in mESCs. Enhancers are arranged in ascending order based on their input-normalized H3K27ac ChIP-Seq intensities. Super-enhancers are defined as the population of enhancers above the inflection point (H3K27ac ChIP-Seq intensities cutoff) of the curve. **j**, Tables presenting the enriched transcription factors (TFs) motifs identified on D-SSTEs in mESCs. **k**,**l**, Metagene profiles showing the averaged read density of specific transcription factors (Oct4, Sox2, Nonag, c-Myc, Klf4, Brd4, Med1, Med12, Cdk8, Cdk9, E2f1, and Tcfcp2l1) ChIP-seq data across S-SSTEs, D-SSTEs, and DSEs in mESCs. Regions spanning 5 kb upstream and 5 kb downstream from the center of regulatory elements are shown.

**Extended Data Fig. 10.**
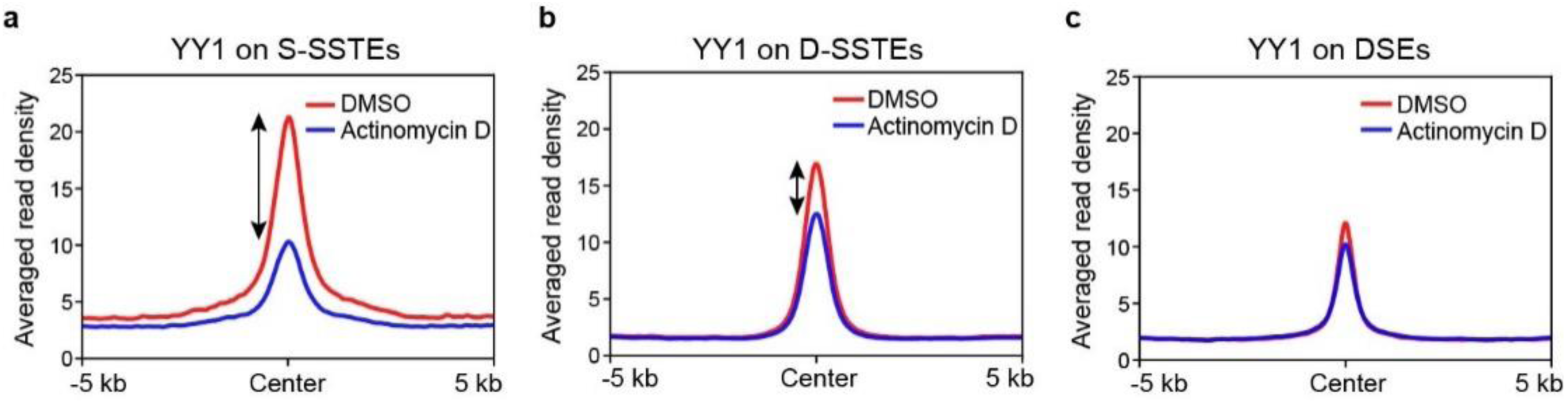
Transcription perturbation by Actinomycin D treatment has varying effects on YY1 binding to different types of SSTEs in mESCs. **a**,**b**,**c**, Metagene profiles showing the averaged read density of YY1 ChIP-seq data obtained from DMSO (red) or Actinomycin D (blue) (transcription inhibition) treated mESCs, across S-SSTEs (**a**), D-SSTEs (**b**), and DSEs (**c**). Regions spanning 5 kb upstream and 5 kb downstream from the center of cis-regulatory elements (CREs) are shown.

**Extended Data Fig. 11.**
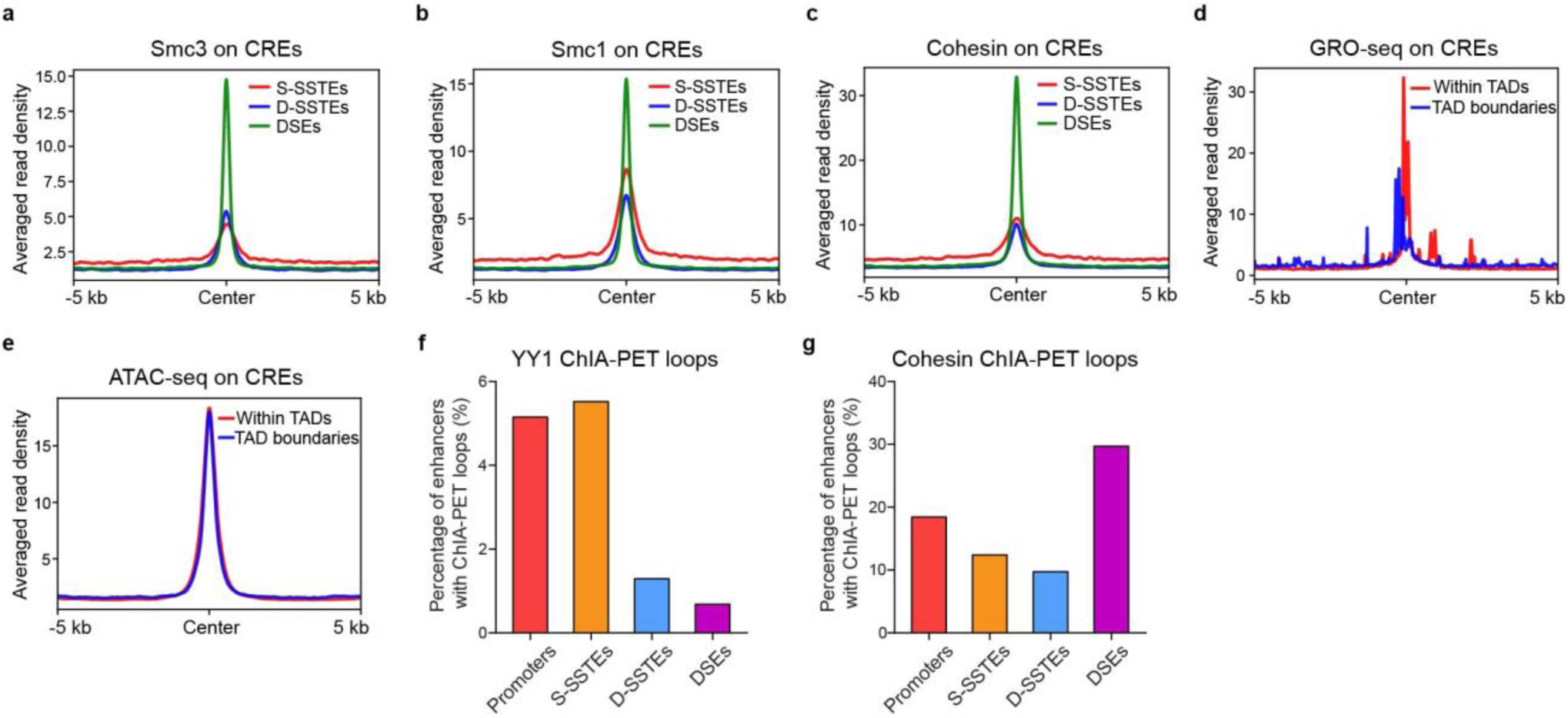
Interaction between the Cohesin complex and associated ChIA-PET loops across different SSTE subtypes in mESCs. **a**,**b**,**c**, Metagene profiles showing the averaged read density of Smc3 (**a**), Smc1 (**b**), and Cohesin (**c**) ChIP-seq data across S-SSTEs, D-SSTEs, and DSEs in mESCs. **d**,**e**, Metagene profiles showing the averaged read density of GRO-seq (**d**) and ATAC-seq (**e**) data across SSTEs within TADs, or at TAD boundaries in mESCs. **f**,**g**, Vertical bar plot illustrating the percentages of promoters, S-SSTEs, D-SSTEs, and DSEs associated with long-range interaction loops defined by YY1 (**f**) and Cohesin (**g**) ChIA-PET data in mESCs.

## Reference

1 Wittkopp, P. J. & Kalay, G. Cis-regulatory elements: molecular mechanisms and evolutionary processes underlying divergence. Nature Reviews Genetics 13, 59–69 (2012).

2 Ong, C.-T. & Corces, V. G. Enhancer function: new insights into the regulation of tissue-specific gene expression. Nature Reviews Genetics 12, 283–293 (2011).

3 Consortium, E. P. An integrated encyclopedia of DNA elements in the human genome. Nature 489, 57 (2012).

4 Jiang, C. & Pugh, B. F. Nucleosome positioning and gene regulation: advances through genomics. Nature Reviews Genetics 10, 161–172 (2009).

5 Schoenfelder, S. & Fraser, P. Long-range enhancer–promoter contacts in gene expression control. Nature Reviews Genetics 20, 437–455 (2019).

6 Sanyal, A., Lajoie, B. R., Jain, G. & Dekker, J. The long-range interaction landscape of gene promoters. Nature 489, 109–113 (2012).

7 Zhang, Y. et al. Chromatin connectivity maps reveal dynamic promoter–enhancer long-range associations. Nature 504, 306–310 (2013).

8 Whalen, S., Truty, R. M. & Pollard, K. S. Enhancer–promoter interactions are encoded by complex genomic signatures on looping chromatin. Nature genetics 48, 488–496 (2016).

9 Core, L. J. et al. Analysis of nascent RNA identifies a unified architecture of initiation regions at mammalian promoters and enhancers. Nature genetics 46, 1311–1320 (2014).

10 Lam, M. T., Li, W., Rosenfeld, M. G. & Glass, C. K. Enhancer RNAs and regulated transcriptional programs. Trends in biochemical sciences 39, 170–182 (2014).

11 Shlyueva, D., Stampfel, G. & Stark, A. Transcriptional enhancers: from properties to genome-wide predictions. Nature Reviews Genetics 15, 272–286 (2014).

12 Li, W., Notani, D. & Rosenfeld, M. G. Enhancers as non-coding RNA transcription units: recent insights and future perspectives. Nature Reviews Genetics 17, 207–223 (2016).

13 Li, W. et al. Functional roles of enhancer RNAs for oestrogen-dependent transcriptional activation. Nature 498, 516–520 (2013).

14 Andersson, R. & Sandelin, A. Determinants of enhancer and promoter activities of regulatory elements. Nature Reviews Genetics 21, 71–87 (2020).

15 Natoli, G. & Andrau, J.-C. Noncoding transcription at enhancers: general principles and functional models. Annual review of genetics 46, 1–19 (2012).

16 Boyle, A. P. et al. High-resolution mapping and characterization of open chromatin across the genome. Cell 132, 311–322 (2008).

17 Creyghton, M. P. et al. Histone H3K27ac separates active from poised enhancers and predicts developmental state. Proceedings of the National Academy of Sciences 107, 21931–21936 (2010).

18 Calo, E. & Wysocka, J. Modification of enhancer chromatin: what, how, and why? Molecular cell 49, 825–837 (2013).

19 Heintzman, N. D. et al. Histone modifications at human enhancers reflect global cell-type-specific gene expression. Nature 459, 108–112 (2009).

20 Yao, L. et al. A comparison of experimental assays and analytical methods for genome-wide identification of active enhancers. Nature biotechnology 40, 1056–1065 (2022).

21 Danko, C. G. et al. Identification of active transcriptional regulatory elements from GRO-seq data. Nature methods 12, 433–438 (2015).

22 Core, L. J., Waterfall, J. J. & Lis, J. T. Nascent RNA sequencing reveals widespread pausing and divergent initiation at human promoters. Science 322, 1845–1848 (2008).

23 Mahat, D. B. et al. Base-pair-resolution genome-wide mapping of active RNA polymerases using precision nuclear run-on (PRO-seq). Nature protocols 11, 1455–1476 (2016).

24 Kwak, H., Fuda, N. J., Core, L. J. & Lis, J. T. Precise maps of RNA polymerase reveal how promoters direct initiation and pausing. Science 339, 950–953 (2013).

25 Kanamori-Katayama, M. et al. Unamplified cap analysis of gene expression on a single-molecule sequencer. Genome research 21, 1150–1159 (2011).

26 Hirabayashi, S. et al. NET-CAGE characterizes the dynamics and topology of human transcribed cis-regulatory elements. Nature genetics 51, 1369–1379 (2019).

27 Windhager, L. et al. Ultrashort and progressive 4sU-tagging reveals key characteristics of RNA processing at nucleotide resolution. Genome research 22, 2031–2042 (2012).

28 Nojima, T. et al. Mammalian NET-seq reveals genome-wide nascent transcription coupled to RNA processing. Cell 161, 526–540 (2015).

29 Buenrostro, J. D., Giresi, P. G., Zaba, L. C., Chang, H. Y. & Greenleaf, W. J. Transposition of native chromatin for fast and sensitive epigenomic profiling of open chromatin, DNA-binding proteins and nucleosome position. Nature methods 10, 1213–1218 (2013).

30 Wu, T., Lyu, R., You, Q. & He, C. Kethoxal-assisted single-stranded DNA sequencing captures global transcription dynamics and enhancer activity in situ. Nature methods 17, 515–523 (2020).

31 Lyu, R. et al. KAS-seq: genome-wide sequencing of single-stranded DNA by N3-kethoxal–assisted labeling. Nature protocols 17, 402–420 (2022).

32 Chen, X. et al. Structural visualization of transcription initiation in action. Science 382, eadi5120 (2023).

33 Wu, T., Lyu, R. & He, C. spKAS-seq reveals R-loop dynamics using low-input materials by detecting single-stranded DNA with strand specificity. Science Advances 8, eabq2166 (2022).

34 Xu, C. et al. R-loop-dependent promoter-proximal termination ensures genome stability. Nature 621, 610–619 (2023).

35 Dou, X. et al. RBFOX2 recognizes N 6-methyladenosine to suppress transcription and block myeloid leukaemia differentiation. Nature cell biology 25, 1359–1368 (2023).

36 Espah Borujeni, A., Zhang, J., Doosthosseini, H., Nielsen, A. A. & Voigt, C. A. Genetic circuit characterization by inferring RNA polymerase movement and ribosome usage. Nature Communications 11, 5001 (2020).

37 Sun, S. et al. Znhit1 controls meiotic initiation in male germ cells by coordinating with Stra8 to activate meiotic gene expression. Developmental cell 57, 901–913. e904 (2022).

38 Fan, H. et al. Trans-vaccenic acid reprograms CD8+ T cells and anti-tumour immunity. Nature 623, 1034–1043 (2023).

39 Grandi, F. C., Modi, H., Kampman, L. & Corces, M. R. Chromatin accessibility profiling by ATAC-seq. Nature protocols 17, 1518–1552 (2022).

40 Kiani, K., Sanford, E. M., Goyal, Y. & Raj, A. Changes in chromatin accessibility are not concordant with transcriptional changes for single-factor perturbations. Molecular Systems Biology 18, e10979 (2022).

41 Mao, X. & Zhao, S. Neuronal differentiation from mouse embryonic stem cells in vitro. JoVE (Journal of Visualized Experiments), e61190 (2020).

42 Kilchert, C., Wittmann, S. & Vasiljeva, L. The regulation and functions of the nuclear RNA exosome complex. Nature Reviews Molecular Cell Biology 17, 227–239 (2016).

43 Lubas, M. et al. Interaction profiling identifies the human nuclear exosome targeting complex. Molecular cell 43, 624–637 (2011).

44 Wu, Y. et al. Nuclear exosome targeting complex core factor Zcchc8 regulates the degradation of LINE1 RNA in early embryos and embryonic stem cells. Cell reports 29, 2461–2472. e2466 (2019).

45 Varlakhanova, N. V. et al. Myc maintains embryonic stem cell pluripotency and self-renewal. Differentiation 80, 9–19 (2010).

46 Filipczyk, A. et al. Network plasticity of pluripotency transcription factors in embryonic stem cells. Nature cell biology 17, 1235–1246 (2015).

47 Sigova, A. A. et al. Transcription factor trapping by RNA in gene regulatory elements. Science 350, 978–981 (2015).

48 Beagan, J. A. & Phillips-Cremins, J. E. On the existence and functionality of topologically associating domains. Nature genetics 52, 8–16 (2020).

49 Dixon, J. R. et al. Topological domains in mammalian genomes identified by analysis of chromatin interactions. Nature 485, 376–380 (2012).

50 Handoko, L. et al. CTCF-mediated functional chromatin interactome in pluripotent cells. Nature genetics 43, 630–638 (2011).

51 Ma, S. et al. Chromatin potential identified by shared single-cell profiling of RNA and chromatin. Cell 183, 1103–1116. e1120 (2020).

52 Corces, M. R. et al. An improved ATAC-seq protocol reduces background and enables interrogation of frozen tissues. Nature methods 14, 959–962 (2017).

53 Li, Y. et al. An optimized method for neuronal differentiation of embryonic stem cells in vitro. Journal of Neuroscience Methods 330, 108486 (2020).

54 Krueger, F. Trim Galore!: A wrapper around Cutadapt and FastQC to consistently apply adapter and quality trimming to FastQ files, with extra functionality for RRBS data. Babraham Institute (2015).

55 Langmead, B. & Salzberg, S. L. Fast gapped-read alignment with Bowtie 2. Nature methods 9, 357–359 (2012).

56 Li, H. et al. The sequence alignment/map format and SAMtools. bioinformatics 25, 2078–2079 (2009).

57 Lyu, R. et al. KAS-Analyzer: a novel computational framework for exploring KAS-seq data. Bioinformatics Advances 3, vbad121 (2023).

58 Stovner, E. B. & Sætrom, P. epic2 efficiently finds diffuse domains in ChIP-seq data. Bioinformatics 35, 4392–4393 (2019).

59 Ramírez, F. et al. deepTools2: a next generation web server for deep-sequencing data analysis. Nucleic acids research 44, W160 (2016).

60 Kuhn, R. M., Haussler, D. & Kent, W. J. The UCSC genome browser and associated tools. Briefings in bioinformatics 14, 144–161 (2013).

61 Quinlan, A. R. & Hall, I. M. BEDTools: a flexible suite of utilities for comparing genomic features. Bioinformatics 26, 841–842 (2010).

62 Gaspar, J. M. Improved peak-calling with MACS2. BioRxiv, 496521 (2018).

63 Ou, J. et al. ATACseqQC: a Bioconductor package for post-alignment quality assessment of ATAC-seq data. BMC genomics 19, 1–13 (2018).

64 Durand, N. C. et al. Juicer provides a one-click system for analyzing loop-resolution Hi-C experiments. Cell systems 3, 95–98 (2016).

65 Li, H. & Durbin, R. Fast and accurate short read alignment with Burrows–Wheeler transform. bioinformatics 25, 1754–1760 (2009).

66 Heinz, S. et al. Simple combinations of lineage-determining transcription factors prime cis-regulatory elements required for macrophage and B cell identities. Molecular cell 38, 576–589 (2010).

67 McLean, C. Y. et al. GREAT improves functional interpretation of cis-regulatory regions. Nature biotechnology 28, 495–501 (2010).

